# Reconstitution of human adrenocortical specification and steroidogenesis using induced pluripotent stem cells

**DOI:** 10.1101/2022.10.31.514433

**Authors:** Yuka Sakata, Keren Cheng, Michinori Mayama, Yasunari Seita, Andrea J. Detlefsen, Clementina A. Mesaros, Trevor M. Penning, Kyosuke Shishikura, Wenli Yang, Richard J. Auchus, Jerome F. Strauss, Kotaro Sasaki

## Abstract

The mechanisms leading to adrenal cortex development and steroid synthesis in humans remain poorly understood due to the paucity of model systems. Herein, we faithfully recapitulate human fetal adrenal cortex specification processes through stepwise induction of human induced pluripotent stem cells through posterior intermediate mesoderm-like and adrenal progenitor-like states to ultimately generate fetal zone adrenal cortex-like cells (FZLCs), as evidenced by histomorphological, ultrastructural, and transcriptome features and adrenocorticotropic hormone (ACTH)-independent Δ5 steroid biosynthesis. Furthermore, FZLC generation is promoted by SHH and inhibited by NOTCH, ACTIVIN and WNT signaling, and that steroid synthesis is amplified by ACTH/PKA signaling and blocked by inhibitors of Δ5 steroid synthesis enzymes. Finally, NR5A1 promotes FZLC survival and steroidogenesis. Together, these findings provide a framework for understanding and reconstituting human adrenocortical development in vitro paving the way for cell-based therapies of adrenal insufficiency.

## Introduction

The adrenal cortex is the major endocrine hub for steroid hormone production, and thus regulates a wide variety of critical physiologic functions vital to human life, including immune and stress responses, sexual maturation, and electrolyte balance. In fetal life, the adrenal cortex also facilitates maturation of various organ systems and is essential for proper functioning of the fetoplacental unit, which plays a role in pregnancy maintenance^1,2^. In addition, successful development of the fetal adrenals is required for establishment of a functional adrenal cortex postnatally^1,3^. Accordingly, genetic defects affecting fetal adrenal development (e.g., *NR5A1*, *WNT4* mutations) or steroid biosynthesis result in primary adrenal insufficiency (PAI), an underdiagnosed but potentially life-threatening disorder necessitating life-long hormone replacement therapy^4^. As there are currently limited therapeutic or preventative measures for PAI, understanding the genetic and signaling requirements for normal fetal adrenal development and initiation of steroidogenesis is of paramount significance for the accurate PAI diagnosis and development of targeted therapeutic interventions.

Our current understanding of human fetal adrenal development has been hampered by the marked differences between human and rodent adrenal development, which limits the translational potential of previous studies using rodent models. For example, adrenal androgens (e.g., DHEA, DHEA-S), the most abundant steroids produced by human fetal adrenals, are only minimally synthesized in rodent fetal adrenals^2,5,6^. In addition, ex vivo culture of human fetal adrenal tissues/cells also suffers from sample-to-sample variability and bioethical constraints^7–11^. Such findings highlight the critical need to develop more suitable models to understand the genetic and epigenetic mechanisms regulating human adrenocortical development and adrenal steroidogenesis.

In the human fetus, organogenesis of the adrenal cortex starts with fate specification of the NR5A1^+^GATA4^−^ adrenogenic coelomic epithelium (AdCE) within the WT1^+^ coelomic epithelium at 3-4 week post fertilization (wpf)^12^. These cells subsequently undergo dorsomedial migration and form a condensed blastematous structure, referred to as the adrenal primordium (AP), which is followed by establishment of two distinct zonal structures, the peripherally located definitive zone (DZ) with putative stem cell/progenitor potential, and the centrally located fetal zone (FZ) with steroidogenic potential by 8 wpf^12–14^. Our single cell RNA-seq analysis of human fetal adrenals has recently revealed the cellular and transcriptional dynamics during specification of adrenocortical lineages^12^. However, the underlying gene regulatory mechanisms governing human adrenocortical development and concomitant steroidogenesis remain unknown.

To overcome these limitations, and establish alternative to the use of human fetal tissue, we explored the possibility of directing human adrenocortical lineages from human induced pluripotent stem cells (hiPSCs) under defined conditions. Using this strategy in multiple hiPSC lines, we observed a robust induction of steroidogenic fetal adrenocortical cells, which we utilized to interrogate human adrenocortical development and steroidogenic function with pharmacologic, genetic and epigenetic approaches.

## Results

### Induction of early adrenocortical lineage through posterior intermediate mesoderm-like cells derived from human iPSCs

Based on our scRNA-seq analyses of human adrenocortical development, we set out to determine if FZLCs could be established from human iPSCs through stepwise induction of 1) T^+^ nascent mesoderm, 2) WT1^+^ posterior intermediate mesoderm (PIM), and 3) NR5A1^+^ adrenocortical progenitors (i.e., AdCE and AP) to finally derive FZLCs. To visualize the stepwise induction process, we generated human iPSCs bearing WT1-p2A-EGFP (WG) (turned on as cells transition to PIM) and NR5A1-p2A-tdTomato (NT) (turned on as cells transition into adrenocortical progenitors) using a CRISPR/Cas9-guided knock-in approach. One male line had both WG and NT (WGNT SV20-211, herein designated as 211) whereas two female lines possessed NT only (NT 312-2121 and NT 1390G3-2125) (Extended Data Fig. 1a-e). As both kidneys and adrenal cortex originate from PIM^15^, we first exploited a previously established 3D induction platform to derive the metanephric lineage by activating Wnt signaling through high dose CHIR99021 treatment for 7 days^16^. This first step of induction generated transient *T*^+^ nascent mesoderm (Extended Data Fig. 2a). Subsequently, at floating culture day (fl)9, retinoic acid-based induction resulted in generation of PIM-like cells bearing high *WT1* but low *FOXF1* expression; the latter representing a marker of the lateral plate mesoderm (Extended Data Fig. 2a-b)^12,15^. Accordingly, further culture of aggregates until fl12 with low CHIR99021 and high FGF9, a potent nephrogenic inducer^16,17^, resulted in *SIX2*-, *PAX2*- and WG-expressing metanephric mesenchyme, consistent with the previous study (Extended Data Fig. 2a-b). As this condition did not activate NR5A1, a key marker of adrenocortical fate (Extended Data Fig. 2a-b)^12^, we next attempted to modify conditions to redirect the lineage towards the adrenal fate using NT fluorescence to monitor our success. Notably, optimization of the initial BMP4 concentration and switching from the original DMEM-based medium supplemented with B27 (DB27 medium) to α-MEM-based medium supplemented with 10% Knockout Serum Replacement (KSR) (MK10 medium) not only increased the induction efficiency of WG, but downregulated metanephric mesenchyme markers (i.e., *SIX2*, *PAX2*) (Extended Data Fig. 2c-d). Moreover, while *NR5A1* remained undetectable, these conditions facilitated *OSR2* expression, one of the earliest markers of adrenocortical fate (Extended Data Fig. 2d)^12,15^, suggesting that it redirected the lineage from the metanephric mesenchyme and might provide the permissive condition for adrenal fate specification. Using these conditions, we explored modulation of factors that might potentially affect adrenal or kidney development. Because CHIR99021, FGF9 and Activin A reportedly enhance nephrogenic fate^16,18^, we first tested inhibitors of these factors. Although antagonism of canonical Wnt and FGF9 by IWR1 and SU-5402, respectively, slightly upregulated NT expression (Extended Data Fig. 2e), addition of SB431542 (NODAL/Activin inhibitor) (fl7-22) together with low dose BMP4 (fl10-15) markedly upregulated NT expression at fl14 and 22 (Fig. 1a). We have previously demonstrated that human fetal adrenal cortex strongly expresses *DLK1*, a membrane bound protein that negatively regulates NOTCH signaling^12,15^. Moreover, Sonic Hedgehog (SHH) is thought to be critical for adrenocortical development, and downstream genes of SHH are highly expressed in human fetal adrenal cortex^12,19–21^. This motivated us to test the effect of DAPT (NOTCH inhibitor) and SHH on NT expression. Notably, DAPT and SHH in combination further enhanced both NT and steroidogenic gene expression at fl22 (Fig. 1a, Extended Data Fig. 2f-j). Notably, this combination also suppressed expression of WT1 and the podocyte marker NPHS2, and lowered the frequency of WG^bright+^ cells, an apparent off-target nephrogenic population (Extended Data Fig. 2h, j). As this condition (SHH: fl7-22; DAPT: fl10-22) combined with early addition of SB431542 (fl5-22) outperformed that of late addition of SB431542 (fl7-22) (Extended Data Fig. 3a), we employed this new modification for subsequent cultures (SHH: fl7-22; DAPT: fl10-22; SB431542: fl5-22). As we now had NT induction, we went back to reevaluate the role of Wnt signaling at fl10-22 in NT induction. Accordingly, we confirmed that IWR1 outperformed CHIR99021 or no-treatment, further suggesting that Wnt activation (fl1-10) followed by inhibition (fl10-22) supports the induction of NT^+^ cells (Extended Data Fig. 3b). Of note, although the culture condition from fl15 onwards described above essentially replicated the preceding condition from fl10 (BISSSD medium), overall concentrations of growth factors/inhibitors were reduced and BMP4 was omitted (ISSSD medium). This, however, did not affect NT induction efficiency (Extended Data Fig. 3c). In this final induction scheme, we confirmed that the initial dose of BMP4 at fl0-1 affects overall induction efficiency at fl22 (Extended Data Fig. 3d).

**Fig. 1.**
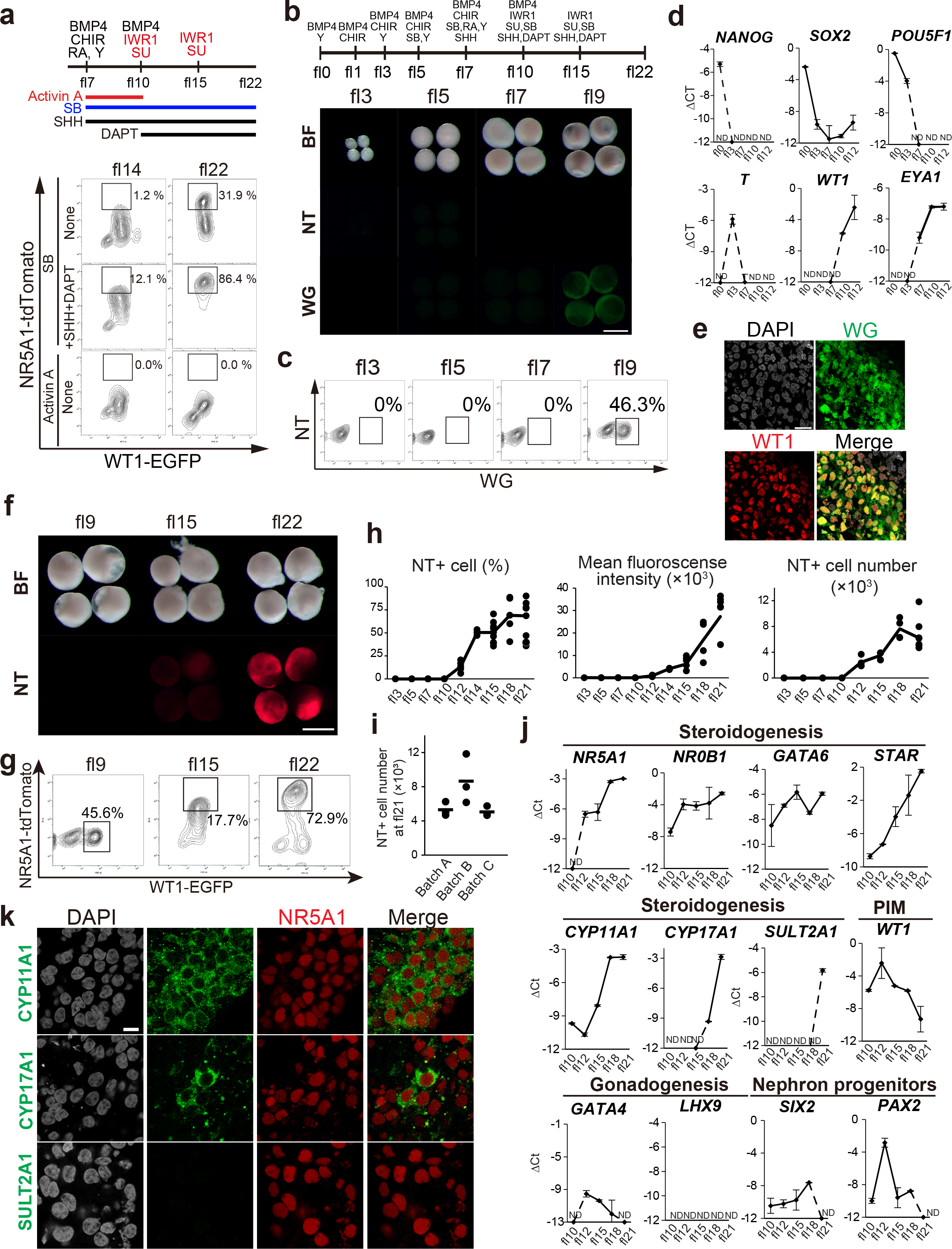
Induction of early adrenocortical lineage through the posterior intermediate mesoderm (PIM)-like cells from human iPSCs. (**a**) Schematic showing several key factors (depicted as horizontal lines) for induction of NT^+^ cells in WGNT SV20-211 (211) hiPSCs (top). Cells were treated with 10 ng/ml ACTIVIN A (fl7-10) or 30 μM SB (in the presence or absence of 50 ng/ml SHH [fl7-22]/ 10 μM DAPT [fl10-22]). Other than these factors, factors depicted above the timeline (fl7-22) and those in the scheme in (**b**) (fl0-7) were used for all culture conditions. BMP4, human BMP4; CHIR, CHIR99021; RA, retinoic acid; SB, SB-431542; SHH, sonic hedgehog; SU, SU-5402; Y, Y-27632. (bottom) Fluorescence-activated cell sorting (FACS) analysis at floating culture day 14 (fl14) and fl22 of the indicated culture condition. (**b**) Schematic depicting the finalized induction scheme (top). Bright-field (BF) and fluorescence images of aggregates for WT1-EGFP (WG, green) and NR5A1-tdTomato (NT, red) at indicated stages (middle). Bar, 500 μm. (**c**) FACS analysis of aggregates at indicated stages. (d) qPCR analysis of expression of key genes in cDNA generated from bulk aggregates derived from 211 hiPSCs at the indicated time point during PIM induction. ΔCt values were calculated by subtracting the raw Ct values of each gene (mean value of two biological replicates) from the averaged Ct values of the housekeeping genes ARBP and PPIA. (**e**) IF images of the fl10 aggregate for WT1-EGFP (green), WT1 (red) and DAPI (white) with their merge. Bar, 20 μm. (**f**) Bright field (BF) and fluorescence (NT, red) images of fl9, fl15 and fl22 aggregates. Bar, 500 μm. (**g**) FACS analysis of WGNT expression in (**f**). (**h**) Percentage of NT^+^ cells, mean fluorescence intensity of NT, and NT^+^ cell number per aggregate during floating culture as assessed by FACS. Cells were derived from 211 hiPSCs. (**i**) NT^+^ cell number per aggregate at fl21 in three different batches. (**j**) qPCR analysis of expression of key genes in cDNA generated from bulk aggregates at the indicated time point. Cells were derived from 211 hiPSCs. (**k**) IF images of fl22 aggregates for CYP11A1, CYP17A1, or SULT2A1 (green), stained with NR5A1 (red) and DAPI (white). Merged images are shown on the right. Bar, 10 μm.

Remarkably, similar to the original conditions, this new multi-step induction first established WT1^+^ PIM-like cells (PIMLCs) at fl9-10, in which PIM markers were upregulated while pluripotency associated markers were downregulated (Fig. 1b-e). Thereafter, aggregates progressively upregulated NT and adrenocortical markers, *NR5A1* and *NR0B1* from fl12 onwards while gradually decreasing WT1 from fl15 onwards, consistent with their commitment to early adrenocortical lineages (Fig. 1f-j)^12^. Importantly, neither gonadal markers (*GATA4*, *LHX9*) nor kidney markers (*PAX2*, *SIX2*) were detectable by fl21, suggesting that cells were directed towards the adrenal cortex but not developmentally related lineages (Fig. 1j). We also noted that cells progressively acquired steroidogenic gene expression including *STAR*, *CYP11A1*, which was followed by lower level upregulation of *CYP17A1* and *SULT2A1* after fl18 (Fig. 1j). Accordingly, IF revealed that CYP11A1 was diffusely expressed in NR5A1^+^ cells whereas CYP17A1 was expressed weakly and only in a few scattered cells and SULT2A1 was undetectable, suggesting that cells were still immature and had not yet acquired the full steroidogenic activity as seen in fetal adrenal cortex in vivo (Fig. 1k).

Similar to in vivo human fetal adrenal cortex, we found that DLK1 is highly expressed in NT^+^ but rarely in NT^−^ cells (Extended Data Fig. 3e-f). Notably, this allowed us to FACS-sort adrenocortical lineages induced from a hiPSC line that did not bear NT alleles (Penn067i-312-1, herein designated as 312); the sorted DLK1^bright+^ cells from 312 hiPSCs subjected to our induction protocol expressed *NR5A1*, *NR0B1*, *STAR* and *CYP17A1* at a level equivalent to or slightly higher than sorted NT^+^ cells from NT 312-2121 hiPSCs, supporting the utility of DLK1 as a cell surface marker of early adrenocortical lineages (Extended Data Fig. 3g).

### Induction of fetal-zone adrenocortical-like cells (FZLCs) through air-liquid interface culture

We noted that culturing aggregates beyond fl22 resulted in increased central necrosis, likely due to limited oxygen and/or nutrient diffusion (Extended Data Fig. 4a). To enhance survival and maturation of aggregates, we transferred them onto the collagen-type I-coated PET membranes for air-liquid interface (ALI) culture (Fig. 2a-c, Extended Data Fig. 4b). Notably, we found that use of ISSD medium (low dose IWR1, SU, SHH, DAPT) outperformed basal MK15 medium for promoting survival of NT^+^ cells during ALI culture (Extended Data Fig. 4c-d). Moreover, ALI culture resulted in progressive upregulation of genes encoding proteins involved in steroidogenesis (i.e., *CYP11A1*, *CYP17A1*, *SULT2A1*, *CYP11B1*), cofactors (e.g., *STAR*, *POR*, *FDX1*, *FDXR*, *PAPSS2*) and other genes related to adrenal steroidogenesis while maintaining NT and NR5A1 expression (Fig. 2b-c, Extended Data Fig. 4e-g). Transcription factors such as *NR0B1*, *OSR2* and *GATA6* were expressed throughout the ALI culture. Metanephric (e.g., *SIX2*, *PAX2*) and gonadal genes (e.g., *GATA4* and *LHX9*) were not expressed, suggesting that the culture conditions did not support these cell types (Extended Data Fig. 4f-g). H&E sections of aggregates at ali21-28 revealed tightly packed nests of cells with abundant eosinophilic and granular cytoplasm and occasional cytoplasmic vacuoles, and peripherally located round nuclei, highly reminiscent of the FZ (Fig. 2e, Extended Data Fig. 4h). IF also revealed strong cytoplasmic expression of CYP11A1, CYP17A1, SULT2A1, SOAT1, FDX1, and FDXR, similar to the FZ (Fig. 2f-g, Extended Data Fig. 4i)^12^. ISH further confirmed the expression of *NR5A1*, *NR0B1*, *STAR*, *APOA1* (Extended Data Fig. 4j). The characteristic cellular morphology and immunophenotype was maintained until at least ali62 (Extended Data Fig. 4k). During ALI culture, aggregates increased production of IGF-II, a key growth factor of the fetal adrenal cortex, that is produced by the FZ, but not by the adult adrenal cortex (Fig. 2h)^22^. Ultrastructurally, these cells had abundant mitochondria that exhibited a leaf-like contour with tubular cristae and intervening prominent smooth endoplasmic reticulum. There were also frequent round osmiophilic lipid granules with various electron density (Fig. 2i). Cells with similar immunophenotypic features were successfully induced from various human iPSC cell lines with or without fluorescence reporters (NT 312-2121, Penn067i-312-1) (Extended Data Fig. 4l, m). Together, gene expression, histologic, immunophenotypic, and ultrastructural features were all consistent with the FZ of the prenatal human adrenal cortex, which led us designate these NT^+^ cells as human fetal zone adrenocortical-like cells (FZLCs).

**Fig. 2.**
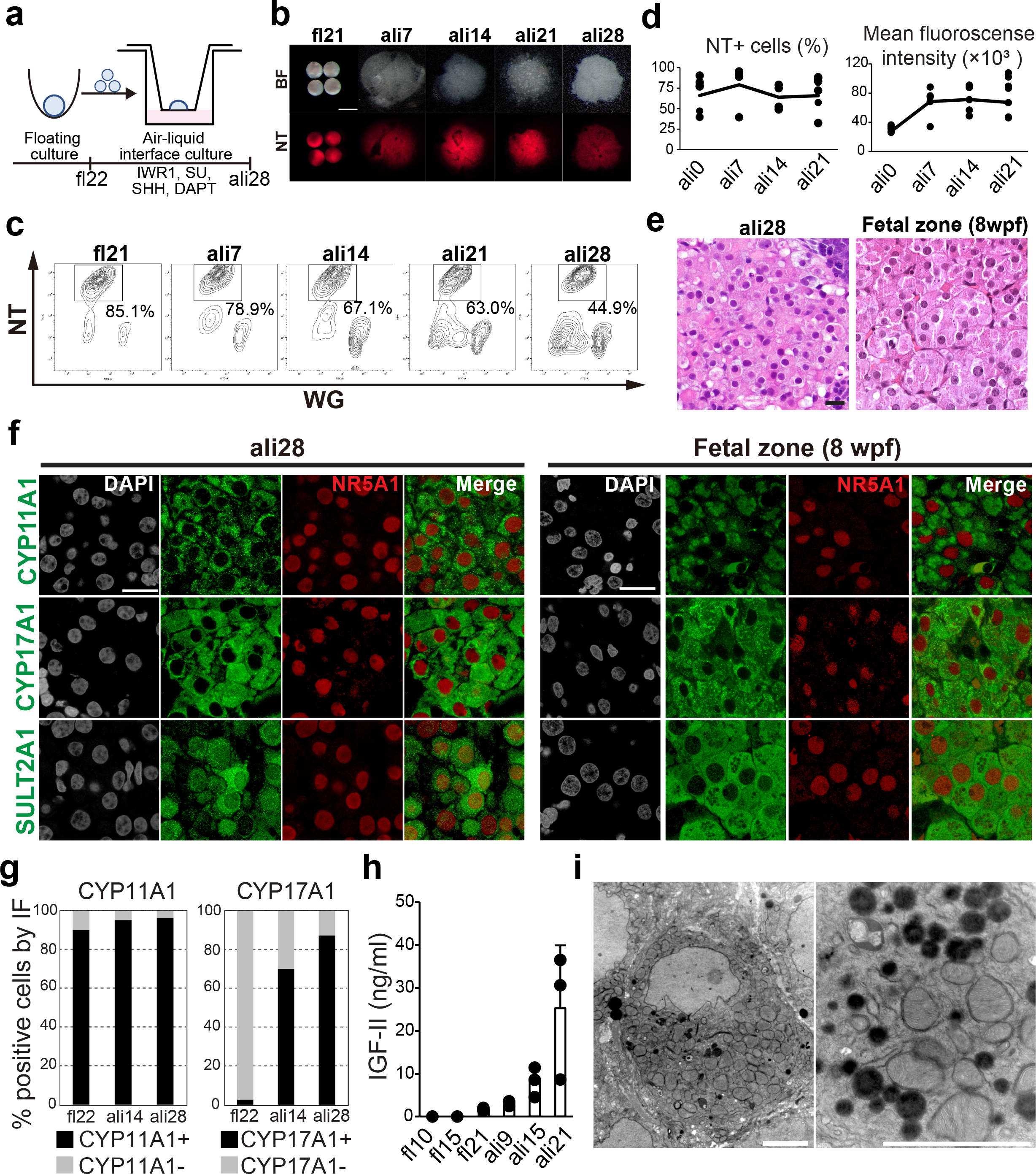
Induction of fetal-zone adrenocortical-like cells (FZLCs) through air-liquid interface culture. (**a**) Schematic illustration of the air-liquid interface (ALI) culture and medium components added during ALI culture. (**b**) BF and fluorescence (NT) images of aggregates at indicated time points. Bar, 500 μm. (**c**) FACS plots of aggregates for WG and NT expression at indicated time points. (**d**) Percentage of NT^+^ cells, the mean fluorescence intensity of NT during ALI culture of 211 iPSC-derived aggregates as assessed by FACS. (**e**) H&E images of the fetal zone (FZ)-like Cells (FZLCs) at ali28 (left) and FZ from a human fetal adrenal gland at 8 wpf (right). Bar, 20 μm. (**f**) IF images of the ali28 (left) and FZ at 8 wpf for CYP11A1, CYP17A1, or SULT2A1 (green), stained with NR5A1 (red) and DAPI (white) and the merged images. (**g**) The IF quantifications of CYP11A1 (left) and CYP17A1 (right) positive cells in aggregates of indicated stages. (**h**) Insulin like growth factor II (IGF-II) production during induction of FZLCs derived from 211 hiPSCs. Culture supernatants 72 h after the last medium change were collected at indicated time points and IGF-II concentration was determined by enzyme-linked immunosorbent assay (ELISA). Means ± standard deviation (n = 3). (**i**) Transmission electron microscope images of the Fetal Zone like cells (FZLCs) in Ali28. Bar, 3 μm.

### Robust Δ5 adrenal steroid biosynthesis from FZLCs

Given the marked upregulation of steroidogenic enzymes in FZLCs, we next measured adrenal steroid production, a key function of the adrenal cortex (Fig. 3a). Consistent with the expression of proteins involved in Δ5 steroid synthesis (e.g., CYP11A1, STAR, CYP17A1 and SULT2A1) (Fig. 2f, Extended Data Fig. 4f, g, i-m), culture supernatant of ali21-28 aggregates contained substantial amounts of Δ5 steroids, dehydroepiandrosterone [DHEA] and its sulfated form [DHEA-S] (Fig. 3b, c). In contrast, most Δ4 steroids were produced in low/negligible levels (e.g., progesterone [prog], 17-OH progesterone [17OHProg], cortisol, aldosterone), except androstenedione (A4), which was produced in moderate levels likely from DHEA through the action of HSD3B2, which was expressed at low but detectable levels (Fig. 3c, Extended Data Fig. 4f). These findings are consistent with the steroid synthesis profile of the human fetal adrenal cortex, which predominantly produces DHEA and DHEA-S^1^. Finally, consistent with the sequential expression of *CYP11A1*, *CYP17A1* and *SULT2A1* (Fig. 1j, Extended Data Fig. 4f), we first detected pregnenolone production at fl21 (Fig. 3d). From ali9 onwards, pregnenolone gradually decreased while DHEA production peaked at ali9 declining thereafter. DHEA-S was also first detected at ali9, but its levels progressively increased until ali21 (Fig. 3d).

**Fig. 3.**
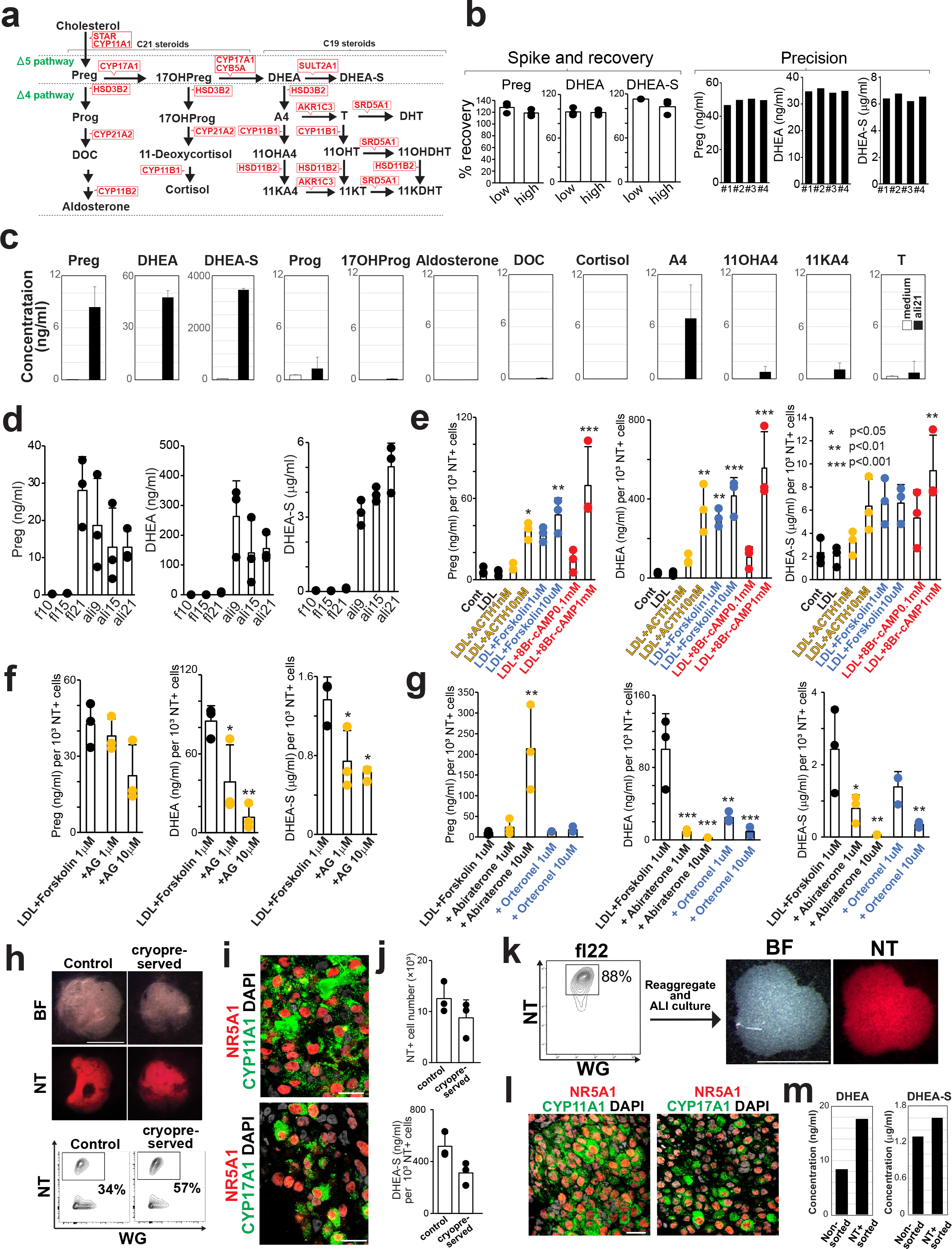
Robust Δ5 adrenal steroid biosynthesis by FZLCs. **(a) Schematic illustration of adrenal steroid synthesis.** Preg, pregnenolone; 17OH Preg, 17α-hydroxypregnenolone; DHEA, dehydroepiandrosterone; DHEA-S, dehydroepiandrosterone-sulfate; Prog, progesterone; 17OH Prog, 17α-Hydroxyprogesterone; A4, androstenedione; T, testosterone; DHT, dihydrotestosterone; DOC, 11-deoxycorticosterone; 11OHA4, 11-hydroxyandrostenedione; 11KA4, 11-ketoandrostenedione; 11OHT, 11-hydroxytestosterone; 11KT, 11-ketotestosterone; 11OHDHT, 11-hydroxydihydrotestosterone; 11KDHT, 11-ketodihydrotestosterone. (**b**) Validation of ELISA assays by spike and recovery and precision measurements using culture medium of al21 aggregates derived from 211 hiPSCs. Low and high concentrations of spiked standard diluent were used for Preg (3.2 ng/ml and 12.8 ng/ml), DHEA (5 ng/ml and 15 ng/ml), and DHEA-S (2.5 μg/ml and 5 μg/ml). Production of Preg, DHEA, DHEA-S was measured using four technical replicates to analyze the precision of the ELISA assay. (**c**) Production of Δ5 and Δ4 steroids in culture media of ali21 aggregates. After a medium change at ali21, aggregates were cultured for additional 3 days for collection of media. As a control, fresh ISSD medium was measured for steroids. ELISA measured Δ5 steroids and LC-MS/MS measured Δ4 steroids. Means ± standard deviation (n = 2). (**d**) Production of Preg, DHEA, and DHEA-S during FZLC induction from 211 hiPSCs. The culture medium was collected at 72 hours and concentrations were determined by ELISA. Means ± standard deviation (n = 3). (**e-g**) Effects of trophic stimulants (ACTH, forskolin, and 8Br-cAMP) (**e**), aminoglutethimide (AG) (**f**), and abiraterone or orteronel (**g**) on the production of Preg, DHEA, and DHEA-S in culture supernatants. Means ± standard deviation (n = 3). Values were normalized with cell numbers. LDL, low density lipoprotein. * p< 0.05, ** p< 0.01, *** p< 0.001 vs control (**e**) and vs LDL + forskolin 1 μM (**f, g**). (**h**) BF and fluorescence images (top) and FACS plots (bottom) of aggregates at ali15 with or without cryopreservation. Aggregates were dissociated at fl21 and cryopreserved before being thawed and reaggregated using 60,000 cells. Bar, 1mm. (**i**) IF images of (**h**) (ali15 after cryopreservation) for NR5A1 (red), CYP11A1, and CYP17A1 (green) merged with DAPI (white). (j) NT^+^ cell number (per aggregate) of aggregates after cryopreservation at ali15 (top) and DHEA-S production measured in culture medium (bottom). Values were normalized with 10^3^ NT^+^ cell numbers (bottom). Means ± standard deviation (n = 3). (k) FACS plot of aggregates at fl22 (left), from which NT+ cells were FACS-sorted and reaggregated for further ALI culture up to ali28. Bright-field (BF) and fluorescence (NT) images of the ali28 (right). Bar, 500 μm. (**l**) IF images of (**k**) for CYP11A1/CYP17A1 (green) and NR5A1 (red) merged with DAPI (white). Bar, 20 μm. (**m**) DHEA and DHEA-S production in supernatant of non-sorted and NT^+^ sorted cells as in (**k**).

ACTH is a polypeptide hormone secreted by the anterior pituitary gland, which plays a central role in the hypothalamic-pituitary-adrenal (HPA) axis by stimulating cell differentiation and steroidogenesis in the adrenal cortex through PKA/cAMP signaling pathways^2^. Therefore, we next evaluated FZLC responsiveness to ACTH. We found that both ACTH and the synthetic PKA/cAMP agonists, forskolin and 8Br-cAMP enhanced the production of pregnenolone, DHEA and DHEA-S in a dose-dependent manner, suggesting that like FZ, FZLCs are responsive to ACTH (Fig. 3e).

Given that FZLCs faithfully recapitulated Δ5 steroid biosynthesis, we next exploited our novel platform for functional interrogation of the adrenal androgen biosynthesis pathway using pharmacologic inhibitors. CYP11A1 is the enzyme required to generate pregnenolone from cholesterol, the first step of steroid biosynthesis. Accordingly, aminoglutethimide (AG), an inhibitor of CYP11A1, dose-dependently suppressed the production of pregnenolone, DHEA and DHEA-S (Fig. 3f). Likewise, CYP17A1 catalyzes both 17α-hydroxylase activity and 17,20-lyase activity to generate DHEA from pregnenolone. Abiraterone inhibits both activities mediated by CYP17A1, while orteronel is a more selective inhibitor of 17,20-lyase activity^23^. Consistently, both drugs dose-dependently suppressed DHEA and DHEA-S synthesis (Fig. 3g). On the other hand, pregnenolone levels increased upon treatment with abiraterone, but not with orteronel, suggesting that selective inhibition of 17,20-lyase by orteronel likely results in the accumulation of 17α-hydroxypregnenolone (Fig. 3g). Altogether, these data suggest that adrenal androgen biosynthesis occurs through the conventional pathway in FZLCs and that our platform could serve as tractable tool for screening inhibitors of this pathway.

### Freeze-thawing, fractionation of aggregates before ALI culture

Given the protracted time required to obtain FZLCs (total of ~43 days), we next determined if cells at fl22 could be cryopreserved before starting ALI culture, thus facilitating subsequent studies. Aggregates were dissociated then frozen in Cell Banker in liquid nitrogen. After thawing, cells were reaggregated in 96 well low-attachment plates for 1 day in the presence of ISSD medium, then transferred to ALI culture for a subsequent 21 days. Notably, these aggregates maintained strong NT and key steroidogenic enzyme expression (i.e., CYP11A1, CYP17A1 and SULT2A1) (Fig. 3h-j). Accordingly, freeze-thawed aggregates exhibited robust production of DHEA-S albeit at slightly lower levels than those without freeze-thawing/reaggregation (not significant, *p* = 0.059). This suggests that aggregates after floating culture can be cryopreserved for future studies.

To further optimize our culture conditions, we next determined if the generation of off-target cells during ALI culture could be minimized by sorting NT^+^ cells prior to culture initiation. For this, sorted NT^+^ cells from fl22 aggregates were allowed to reaggregate in 96 well low-attachment plates and ALI culture for 21 days. These aggregates also maintained bright NT fluorescence and steroidogenic enzymes and exhibited robust steroid synthesis, suggesting that NT^+^ cells can differentiate into FZLCs that are competent to produce Δ5 adrenal androgens even in the absence of NT^−^ cells (Fig. 3k-m).

### Gene expression dynamics during human adrenal specification in vitro

To capture the global transcriptional dynamics of the FZLC induction process, we performed bulk RNA-seq analyses of key cell types during the induction (Fig. 4a-f, Extended Data Fig. 5a, Supplementary Table 1-2). To focus our analyses on adrenocortical lineages, we FACS-sorted WG^+^ cells from fl9 aggregates and NT^+^ cells from fl14 onwards. Unsupervised hierarchical clustering (UHC) segregated the cells into two large clusters, one with hiPSCs, fl9 and fl14 aggregates and the other with fl22 and ali7-28 aggregates (Fig. 4a). Consistently, principal component analysis (PCA) revealed a directional and progressive transition of transcriptional properties during FZLC induction (Fig. 4b). Concordant with qPCR and IF studies, hiPSCs-to-fl9 transition was characterized by upregulation of PIM markers (e.g., *WT1*, *OSR1* and *EYA1*) and *HOX* genes (e.g., *HOXA1*, *HOXB2*) and accordingly, enriched with GO terms such as “multicellular organism development” (Fig. 4c, Supplementary Table 1). In keeping with NT activation, fl14 aggregates showed upregulation of transcription factors associated with adrenal specification (e.g., *NR5A1*, *NR0B1*, *GATA6*, *RUNX1T1*) ^12,15^. This was followed by progressive upregulation of a number of genes associated with adrenal steroid biosynthesis from fl22 onwards with the enriched GO terms such as “cholesterol metabolic process” (fl14-to-fl22 comparison), “cholesterol biosynthetic process” or “regulation of lipid metabolic process” (fl22-to-ali21-28 comparison) (Fig. 4c, Supplementary Table 1). Notably, expression of PIM markers was maintained until fl14, then progressively downregulated (Fig. 4c-d). Accordingly, fl14 aggregates clustered with fl9 by UHC (Fig. 4a), with DEGs enriched with GO terms such as “pattern specification process” (Extended Data Fig. 5b, Supplementary Table 2) and co-expressed adrenocortical and PIM/coelomic epithelium markers (e.g., *WT1*, *BMP4*, *KRT18*, *KRT19*), similar to AdCE in human embryos (Fig. 4c-d)^12^. Likewise, fl22 aggregates highly expressed markers of AP and to a lesser extent, those of AdCE (Fig. 4d). Accordingly, fl9-to-fl22 changes highly resembled transcriptomic dynamics of PIM/early coelomic epithelium (ECE)-to-AP transition in cynomolgus monkey embryos (Extended Data Fig. 5c-d, Supplementary Table 3)^15^. Finally, to further validate our model, we compared the transcriptomes of in vitro derived cells with that of in vivo. To this end, we generated cDNA for whole fetal adrenal cortex at 9 wpf (Ad) and FACS-sorted MME^−^ adrenocortical cells at 17-19 wpf, that were enriched with FZ (Extended Data Fig. 5e)^12^. These samples were sequenced using the same platform used for in vitro cells during FZLC induction. Remarkably, transcriptomes of FZLCs during ALI culture (ali7-28) juxtaposed to and partially overlapped with those of Ad and FZ in PCA coordinate (Fig. 4e). Consistently, these samples exhibited high correlation coefficients (Fig. 4f), and FZLC bore key FZ markers previously defined by single cell RNA-seq of human fetal samples (Fig. 4d). Altogether, these findings suggest that aggregates progressively acquire FZLC state through developmental processes similar to that of human adrenocortical lineage in vivo.

**Fig. 4.**
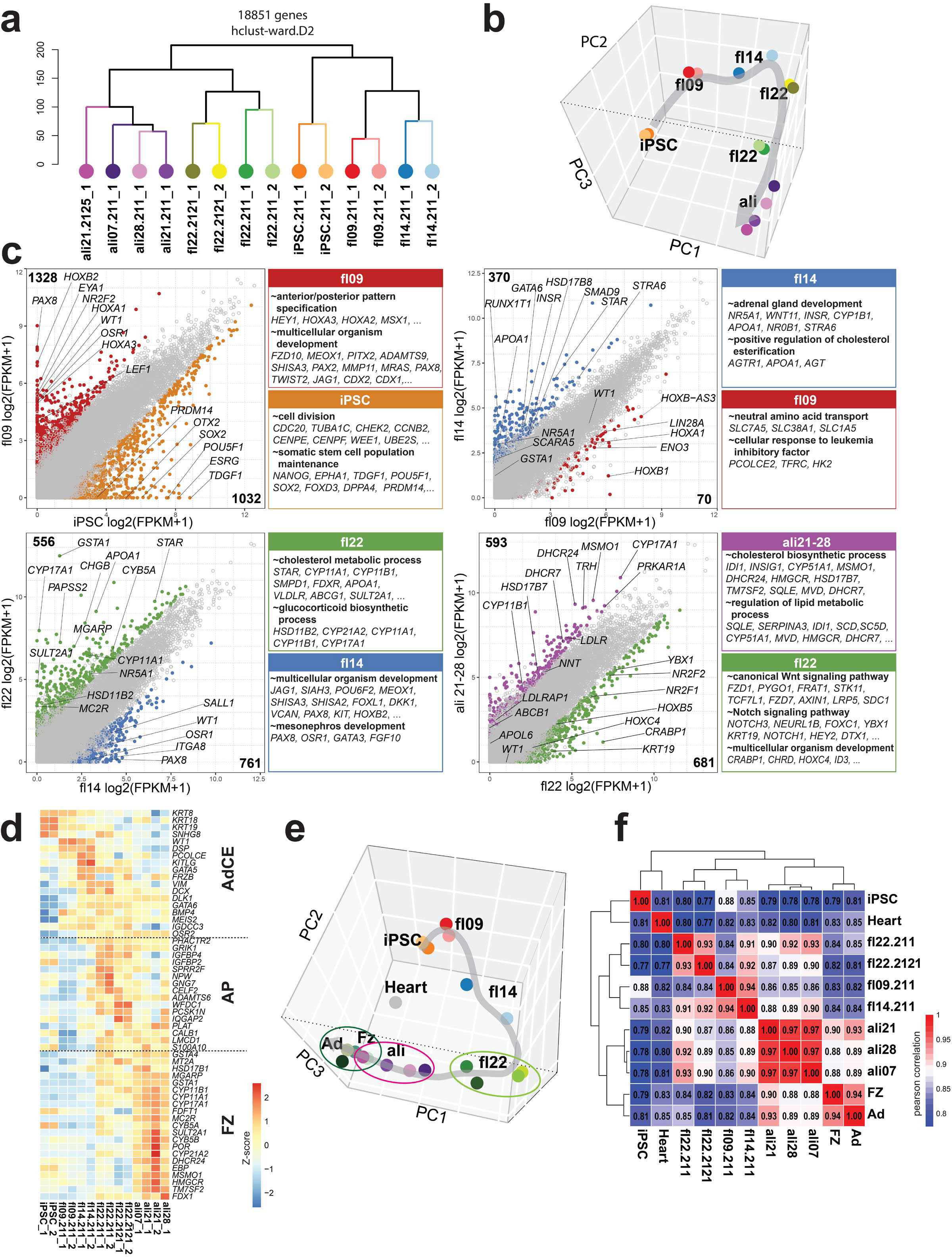
Transcriptome dynamics during FZLC induction. (**a, b**) Unsupervised hierarchical clustering (UHC) (**a**) and principal component analysis (PCA) (**b**) of bulk transcriptomes during FZLC induction; 18851 genes are used for ward.D2 clustering. Cells derived from 211, NT 312-2121 and NT 1390G3-2125 hiPSCs at indicated time points were used. (**c**) Scatter plots comparing the averaged gene expression values between samples at the indicated time points. Differentially expressed genes (DEGs) (four-fold differences, p <0.05, FDR <0.05) are highlighted in colors. Averaged expression values among the samples at the indicated time points are shown. Key genes are annotated. Representative genes and their GO enrichments for DEGs are shown on the right. (**d**) Expression of AdCE, AP and FZ markers in indicated samples as shown by Heatmap. These marker genes were identified as DEGs in previous transcriptome analyses of human fetal samples12. (**e**) PCA of transcriptomes used in (**b**) projected together with human fetal heart, MME-fetal zone adrenocortical cortex (FZ), and whole adrenal glands (Ad). (**f**) Heatmap showing Pearson correlation. Note the high correlation of NT^+^ cells during ali7–28 highly with FZ or Ad in vivo.

### scRNA-seq revealed lineage trajectories of on-target and off-target cell types

To precisely understand the lineage trajectory and transcriptomic dynamics of various cell types derived in vitro at single cell resolution, we performed single cell (sc)RNA-seq using a 10x genomics platform on five samples, of which three were obtained from floating culture (fl19_1, fl19_2, fl22) and two were obtained from ALI culture (ali21, ali 28). All of these samples were FACS-sorted for NT^+^ cells except fl19_2 and ali21, which utilized the WG^+^ fraction or whole live cells, respectively. These cells contained a median of 3572 genes/cell at a mean sequencing depth of 44,499 reads/cell. After QC validation and filtering out low quality cells, 23570 cells remained for downstream analysis (Extended Data Fig. 6a). Transcriptomes of cells in all samples were aggregated and projected onto a UMAP plot after dimension reduction and clustering, which yielded 13 clusters (Extended Data Fig. 6b). All clusters except cluster 13 were successfully annotated based on marker gene expression and differentially expressed genes (Fig. 5a-d, Extended Data Fig. 6c-e, Supplementary Table 4). As expected, we identified *NR5A1*^+^*CYP11A1*^+^*CYP17A1*^+^*STAR*^+^ FZLCs (clusters 3 [AP/early fetal zone like cells] and 6 [fetal zone like cells], consisting primarily of ali21, ali28 aggregates) and their putative progenitor cell types including *WT1*^+^*SFRP2*^+^*SHISHA2*^+^ PIM/CE-like cells (cluster 5) and *NR5A1*^+^*OSR2*^+^*DLK1^+^KRT19*^+^ AdCE-like (clusters 1 and 8, consisting primarily of fl19, 22 aggregates), which aligned along the pseudotime trajectory and actual sample stages (Fig. 5a-d). Along this trajectory, developmental genes or genes related to cell cycle were downregulated (GO terms include “multicellular organism development” and “positive regulation of cell proliferation”), suggestive of terminal differentiation into the FZ adrenocortical lineage (Fig. 5e). In contrast, genes related to steroidogenesis and angiogenesis (GO terms include “cholesterol metabolic process” and “angiogenesis”, respectively) were upregulated along the trajectory, the latter of which might be involved in formation of extensive vascular networks characteristic of the adrenal cortex (Fig. 5e) ^1^. Interestingly, we also noted that previously identified definitive zone markers (*NOV*, *HOPX*) were transiently upregulated in the trajectory although cells expressing these transcripts did not form independent clusters (Extended Data Fig. 6f). In keeping with this observation, IF analyses revealed a few scattered cells that bore DZ markers (NOV, MME) but lower levels of a FZ marker (FDX1), which tended to localize at the periphery of FZLCs. This finding suggests that our platform not only generated FZLCs but also a small fraction of cells resembling DZ-like cells (Extended Data Fig. 6g-k).

**Fig. 5.**
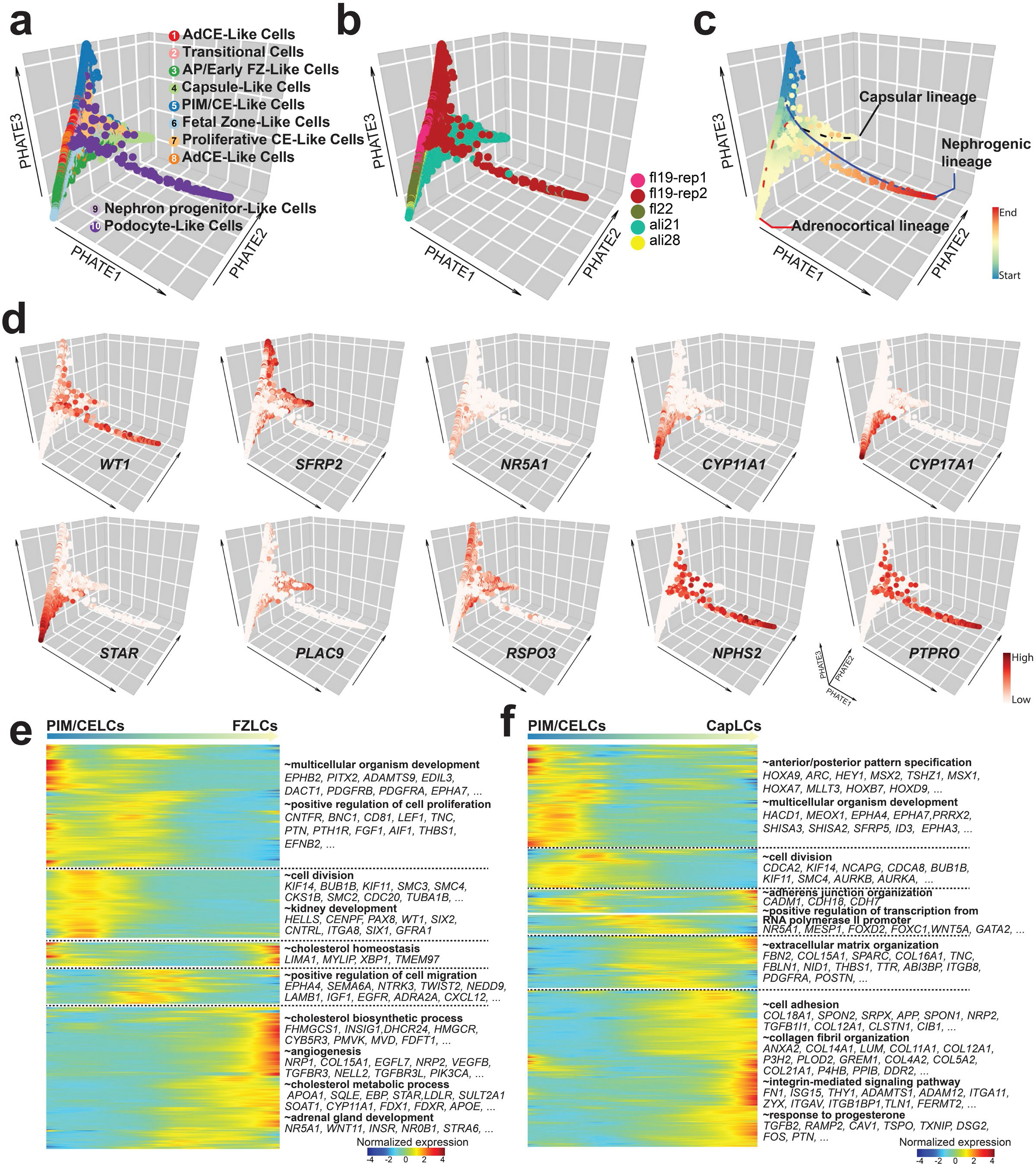
Dynamics of single cell transcriptomes during FZLC induction. (**a-c**) Trajectory analysis of single-cell transcriptomes during FZLC inductions showing segregation of three distinct lineages. Cells are projected to PHATE embedding. Cells are colored according to cell clusters (1: AdCE-like cells, 2: Transition Cells, 3: Fetal zone-like cells, 4: Capsule-like cells 5: PIM/CE-like cells, 6: Fetal Zone-like cells, 7: Proliferative CE-like cells, 8: AdCE-like cells, 9: Nephron progenitor-like cells, 10: Podocyte-like cells). Sample origin (**b**) and pseudotime (**c**) are also projected to the same embedding. (**d**) Expression of key marker genes projected on PHATE embedding. (**e, f**) Heatmap of gene expression along pseudotime trajectory for PIM/CELCs to FZLCs (**e**) or PIM/CELCs to CapLCs (**f**). Genes with similar expression patterns are clustered together, and enriched gene ontology terms are listed alongside the heatmap. The top 2000 variable genes were selected and clustered by UHC for each lineage trajectory. PIM/CELCs, PIM/CE-like cells; CapLCs, capsule-like cells.

Unexpectedly, we also identified minor off-target cell types including *CLDN5*^+^*KDR*^+^ endothelial-like cells and *ASCL1*^+^*GATA3*^+^*PHOX2B*^+^*TH*^+^*DBH*^+^ sympathoadrenal-like cells of uncertain origin (Extended Data Fig. 6b, e)^12^. Notably, trajectory analyses also revealed two additional lineages that appear to project from PIM/CE-like cells, thus sharing their origin with adrenocortical lineages. One of them was the nephrogenic lineage (*SIX2*^+^*PAX2*^+^ nephron progenitor-like cells [cluster 9] and *NPHS2*^+^*PTPRO*^+^ podocyte-like cells [cluster 10]) (Fig. 5a, d, Extended Data Fig. 6b, e), and the other cluster was *RSPO3*^+^*LGALS1*^+^*COL14A1*^+^*PLAC9^+^PDGFRA*^+^ capsule-like cells (CapLCs) (cluster 4), which progressively acquired the characteristic gene expression signature of adrenal capsule cells through a transient proliferative CE-like state (cluster 7) (Fig. 5a, f, Extended Data Fig. 6b, e, 7a-b). Accordingly, CapLCs showed marked transcriptional similarities with the adrenal capsule cells (Cap) (Fig. 5d, Extended Data Fig. 7b-e)^12^. UHC also revealed that CapLCs bore some similarities to PIM/CE/AdCE clusters, expressing shared markers such as *RSPO3* or *SFRP2* (Fig. 5d, Extended Data Fig. 7b-e). Notably, *RSPO3*^+^*PDGFRA*^+^ CapLCs localized at the periphery of the clusters of FZLCs, reminiscent of the capsule in fetal adrenals, although CapLCs were not packed as densely as that of the fetal adrenals (Extended Data Fig. 7f). Together, our data suggest that our inductive scheme not only generate FZLCs through an AdCE-like state but also gives rise to cells with transcriptomic features characteristic of adrenal capsule cells.

### Single cell accessible chromatin landscape of the specifying adrenocortical lineages

NT^+^ cells in fl22 aggregates transcriptionally resembled AP/AdCE of the nascent adrenocortical lineage (Fig. 4d, Extended Data Fig. 5c-d). As these cells emerged from WG^+^ PIM-like cells that were seen in fl9 aggregates (Fig. 1e, g, 4b), this transition recapitulates the adrenocortical lineage specification process. To identify cis-regulatory elements, enhancer regions, and co-accessibility changes during adrenocortical lineage specification, we isolated WG^+^ and NT^+^ cells from fl09 and fl22 aggregates, respectively, and performed single cell Assay for Transposase-Accessible Chromatin sequencing (scATAC-seq). We sequenced a total of 11,365 cells (fl09 WG^+^ cells [8,404 cells] and fl22 NT^+^ cells [2,961 cells]), with 12,148 average median fragments per cell. Overall, we obtained 130,598 unique peaks in fl09 WG^+^ cells, 79,472 unique peaks in fl22 NT^+^ cells, and 120,090 peaks shared between both samples. The majority of peaks were identified in regions characterized as distal elements or introns, which might indicate that they contain enhancer elements (Fig. 6a). The stage-specific peaks showing variability in chromatin accessibility between fl09 and fl22 were enriched in different motifs. Peaks specific for fl09 aggregates revealed motif enrichment related to PIM development, such as SIX1, PAX8, LHX1, LEF1, HOXA1 and CDX2 (Fig. 6b). In contrast, binding motifs for transcription factors potentially involved in adrenocortical development, such as NR5A1, NR2F2, NR2F1, and PBX2, were enriched in peaks uniquely identified in fl22 aggregates. The overlapping peaks, which might represent the housekeeping cis-regulatory elements were enriched with binding motifs for CTCF, which is a constitutive transcription factor binding to enhancers as well as other transcription factors (Fig. 6b).

**Fig. 6.**
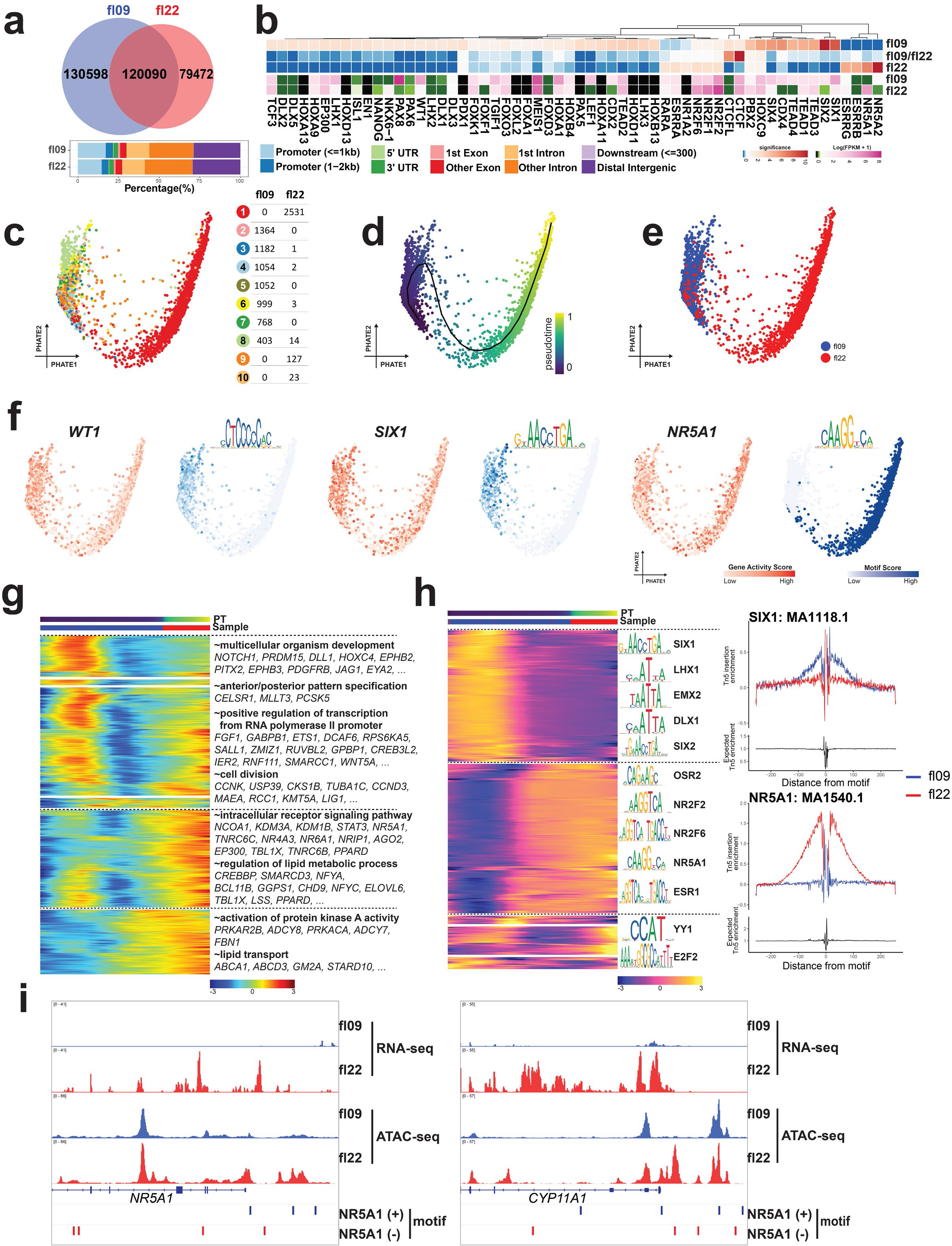
Single cell chromatin accessibility landscape of human adrenocortical specification. (**a**) Genomic annotation of open chromatin regions identified in fl09 and fl22 are categorized in peaks unique to fl09 or fl22 or overlapped between fl09 and fl22. (**b**) The top part of this heatmap shows HOMER motif enrichment analyses using different peaks. Each column is a motif, and colors indicate the motif enrichment significance. The bottom part of this heatmap shows the expression of genes coding for transcription factors correlated with motifs in fl09 and fl22. (**c**) Integration of fl09 and fl22 scATAC-seq. Cells are embedded using PHATE and are clustered using Signac. Ten cell clusters are identified; sample information and cell numbers are listed in the table to the right. (**d**) Trajectory analysis using PHATE and principal trajectory curve fitted on the embedding. Cells are colored according to pseudotime. (**e**) Pseudotime is consistent with the sample information. (**f**) Gene activity scores for WT1, SIX1, NR5A1 and motif enrichment scores for WT1, SIX1, NR5A1 motifs. (**g**) Pseudotime heat-map ordering of gene accessibility across cell differentiation from fl09 to fl22. Gene accessibility is determined by the gene activity score which is generated by counting the read counts per cell in the gene body and promoter region (2000bp upstream of TSS). (**h**) Pseudotime heatmap ordering of chromVAR TF motif across cell differentiation from fl09 to fl22. Right: motif footprint of two representative TF motifs, flanking regions of SIX1 motif is more highly enriched in fl09 than that of fl22. In contrast, flanking regions of the NR5A1 motif is higher enriched in fl22 than that of fl09. There is a typical low-cut frequency in the center of the motif loci. (**i**) Representative genomic browser screenshot of NR5A1 and CYP11A1. In each screenshot, the top two tracks are bulk RNA-seq showing gene expression; the bottom two tracks show the NR5A1 motif loci indicating potential NR5A1 binding sites.

We next determined if transcription factors might be involved in adrenocortical specification by integrating our motif enrichment analyses and transcriptome data. To this end, we first evaluated the dynamics of chromatin accessibility in single cells using latent semantic indexing (LSI) for clustering and PHATE for dimensional reduction (Fig. 6c-e). WG^+^ PIM-like cells at fl09 revealed heterogeneity of open chromatin accessibility as reflected by the increased number of cell clusters (clusters 2-8), whereas chromatin accessibility of fl22 was more homogenous (clusters 1, 9, 10) (Fig. 6c-e). Cellular trajectory and pseudotime inference obtained using stage-specific open chromatin status were concordant with that of the single cell transcriptomes (Fig. 6d-e). We postulated that cell-type specific open chromatin status correlates with cell type-specific gene expression of nearby genes and inferred gene activities based on sums of peaks in proximity of the gene. We found that gene activities of WT1 and SIX1, markers of PIM, diminished along the developmental trajectory (Fig. 6f), consistent with the decline of these transcript levels as assessed by RNA-seq (Fig. 6b). On the other hand, concordant with substantial upregulation of transcript levels (Fig. 6b), chromatin associated with NR5A1, a marker of early adrenocortical lineage, became increasingly open along the trajectory (Fig. 6f). The top 5000 genes based on their variance in accessibility revealed different patterns along pseudotime (Fig. 6g). Genes whose activities were downregulated along pseudotime were those related to early development (enriched with GO terms such as “multicellular organism development” or “anterior/posterior pattern specification”), whereas genes whose activities were upregulated along pseudotime were those related to steroidogenesis or adrenocortical function (enriched with GO terms such as “regulation of lipid metabolic process” or “lipid transport”), which are consistent with our transcriptomic data (Fig. 6g). Moreover, single cell motif enrichment along the pseudotime trajectory revealed footprints of transcription factors related to PIM (e.g., SIX1, LHX1, EMX2) earlier in pseudotime and those related to adrenocortical lineage (e.g., OSR2, NR5A1) later in pseudotime, suggesting that these factors transcriptionally regulate the PIM to adrenocortical cell fate transition through identified cis-elements (Fig. 6h). Accordingly, NR5A1 motifs were frequently identified in intronic and promoter regions of *NR5A1* itself and genes encoding steroidogenic enzymes (e.g., CYP11A1) providing additional insight into its function (Fig. 6i).

### A critical function of NR5A1 in survival and steroidogenesis of FZLCs

In mice, fetal adrenal specification and steroidogenesis is initiated in a dose sensitive manner by the transcription factor, NR5A1^24–27^. Biallelic mutations in murine *Nr5a1* lead to adrenal agenesis and PAI, resulting from defective proliferation and/or increased apoptosis immediately after the formation of the AP^24,25^. However, the functional significance of biallelic *NR5A1* mutations in human fetal adrenal development is unclear. Thus, taking advantage of our FZLC induction platform, we next determined the role of NR5A1 during early adrenocortical development. As simple disruption of *NR5A1* by CRISPR/Cas9 in NR5A1-tdTomato reporter hiPSCs could affect tdTomato expression unpredictably, we established a new NR5A1-knock-in/knock-out hiPSCs in which the DNA binding domain spanning exon 2 and 3 of *NR5A1* was replaced *with* a *2A-tdTomato-polyA* cassette in WG reporter hiPSCs, thus enabling simultaneous gene disruption and visualization of the promoter activities of *NR5A1* (*WGNT-NR5A1^−/−^ SV20-211-19*, herein designated as KIKO) (Extended Data Fig. 1f). Similar to wild-type isogenic 211 hiPSCs, KIKO iPSCs were induced into NT^+^ cells at fl22 with roughly equal induction efficiency, suggesting that *NR5A1* is dispensable for initial lineage specification (Fig. 7a). However, we noted that NT fluorescence intensity, which was already somewhat lower in the mutant line at fl22, was progressively downregulated after ALI culture, suggesting that NR5A1 may autoregulate its promoter activity (Fig. 7a-b), consistent with our scATAC-seq data and a previous report in mice (Fig. 6i)^28^. Remarkably, upon ALI culture, mutant aggregates also exhibited thinning and disintegration, and unlike FZLCs in wild-type aggregates, consisted of partly necrotic cells with karryorrhexis and scanty cytoplasm (Fig. 7b-d). Accordingly, there was markedly diminished recovery of NT^+^ cells and increased cleaved caspase 3^+^ apoptotic cells (Fig. 7c-f). Mutant aggregates also revealed reduced steroidogenic gene and protein expression (Fig. 7g-i) and a consequent dramatic reduction in Δ5 steroid synthesis (Fig. 7j). Together, these findings support the critical role of *NR5A1* in survival and steroidogenesis of human fetal adrenal cortex.

**Fig. 7.**
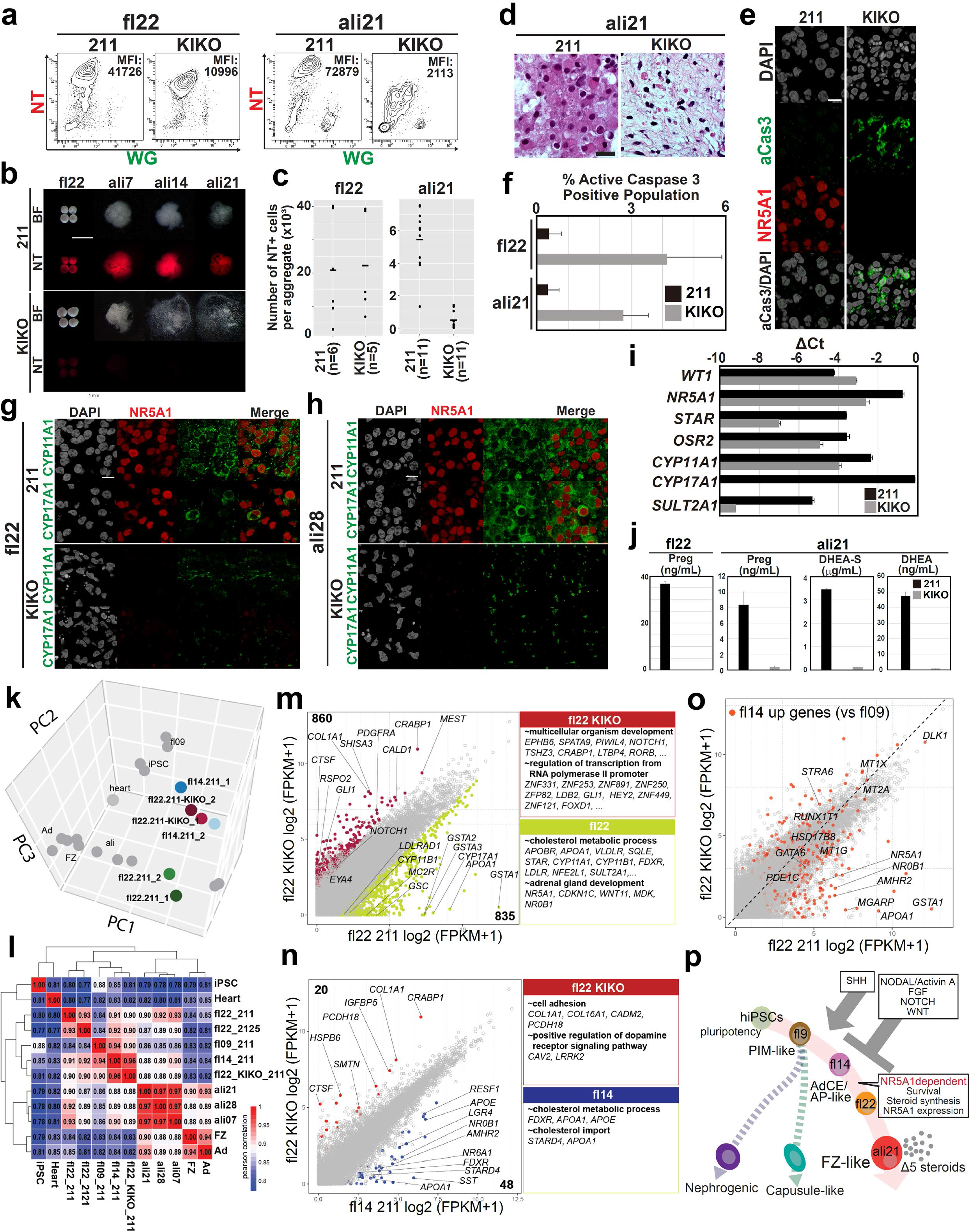
NR5A1 is essential for FZLC survival and steroidogenesis. (**a**) FACS plots of wild-type (211) or NR5A1 mutant (KIKO) aggregates at fl22 (left) and ali21 (right) of induction. MFI, mean fluorescence intensity. (**b**) Bright field (BF) and fluorescence (NT) images of fl22, ali7, ali14 and ali21. Bar, 500 μm. (**c**) Number of NT^+^ cells per aggregate at fl22 and ali21 in wild-type or mutant lines. (**d**) Hematoxylin and eosin images of wild-type or mutant aggregates at ali21. Bar, 20 μm. (**e**) IF images of the wild-type or mutant aggregates at fl22 for active caspase 3 (aCas3) (green), co-stained for NR5A1 (red) and DAPI (white). Merged images for aCas3 and DAPI are shown. Bar,10 μm. (**f**) The quantification of active caspase 3 positive population in (**e**). (**g, h**) IF images of the aggregates at fl22 (**g**) or ali28 (**h**) for CYP11A1 or CYP17A1 (green), co-stained for NR5A1 (red) and DAPI (white). Merged images of NR5A1 and CYP proteins are shown on the right. Bar,10 μm. (**i**) qPCR quantification of gene expression for key marker genes in wild-type and mutant aggregates at fl22. (**j**) Production of Preg, DHEA, and DHEA-S by wild-type (211) or mutant (KIKO) aggregates at indicated time points. (**k**) PCA of transcriptomes of mutant NT^+^ cells at fl22 projected together with those of wild-type NT^+^ cells used in Fig. 4e. Samples at fl14 and fl22 are highlighted in colors. (l) Heatmap of Pearson correlation showing close correlation of mutant (KIKO) aggregates at fl22 to wild-type (211) aggregates at fl14. (**m**) Scatter plot comparison of the averaged values of wild-type (211) and mutant (KIKO) aggregates at fl22. Differentially expressed genes (DEGs) (four-fold differences, p <0.05, FDR <0.05) are highlighted in colors. Red, 860 genes higher in mutants; green, 835 genes higher in wild-type. Key genes are annotated. Representative genes and their GO enrichments for DEGs are shown at the right. (**n**) Scatter plot comparison of the averaged values between wild-type (211) at fl14 and mutant (KIKO) aggregates at fl22. DEGs are highlighted in colors. Representative genes and their GO enrichments for DEGs are shown at the right. (**o**) DEGs upregulated in wild-type fl14 aggregates compared to fl09 aggregates (as defined in Fig. 4c, highlighted in red) projected to a scatter plot used in (**m**). Key genes are annotated. (**p**) Schematic representation showing induction of FZLCs and related lineages from hiPSCs.

To gain further insight into the role of *NR5A1*, we conducted bulk RNA-seq analyses of *NR5A1* mutants. Remarkably, PCA and correlation coefficient heatmaps revealed that mutant NT^+^ cells at fl22 were similar to wild-type NT^+^ cells at fl14, suggesting that NR5A1 is the major driver for the fl14-to-fl22 transition. Accordingly, pairwise comparison of mutant and wild-type NT^+^ cells at fl22 revealed that mutant NT^+^ cells exhibited marked loss of genes related to steroidogenesis, including *CYP17A1*, *CYP11B1*, *APOA1*, *LDLRAD1* and *MC2R*, which were enriched in GO terms, such as “cholesterol metabolic process” or “adrenal gland development” (Fig. 7m, Supplementary Table 5). Like wild-type NT^+^ cells at fl14, mutant NT^+^ cells at fl22 exhibited persistent expression of developmental genes, which were enriched with GO terms such as “multicellular organism development” (Fig. 7m). Although the mutant NT^+^ cells at fl22 were transcriptionally similar to wild-type cells at fl14, a small number of genes were already downregulated in mutants, including *NR6A1*, *NR0B1*, *APOA1*, *FDXR*, *LGR4*, which might represent early genes whose expression is directly driven by NR5A1 (Fig. 7n, Supplementary Table 5). In contrast, some genes that were upregulated in wild-type fl14 cells versus those of fl9 might therefore be involved in adrenal development or steroidogenesis (e.g., *STRA6*, *GATA6*, *DLK1*, *RUNX1T1*, *HSD17B8*, *PDE1C*) were also expressed in mutant cells at fl22, suggesting that initial expression of some early adrenocortical genes is driven by NR5A1-independent mechanisms (Fig. 7o). Together, these data highlight the key role of NR5A1 in early adrenocortical survival and steroidogenesis in humans.

## Discussion

Normal development of the adrenal cortex is essential for fetal and neonatal health, as its disruption leads to PAI and disorders of sex development. Despite its importance, the cellular and molecular mechanisms regulating human adrenocortical development remain unclear, in part due to limited access to fetal adrenal tissues. In this study, we overcome this hurdle by developing an in vitro model system in which adrenocortical lineages can be derived from hiPSCs in a manner that faithfully recapitulates the normal developmental trajectory (Fig. 7p). For this, we first modified a previous nephrogenic induction scheme to maximize the induction of WG^+^ PIM-like cells; then we sought to identify conditions that generate NT^+^ adrenocortical lineages from WG^+^ PIM-like cells in floating culture (Fig. 1). Remarkably, we found that inhibitors, particularly those of NODAL/ACTIVIN or NOTCH signaling, are critical for adrenocortical induction (Fig. 7p). We also found that SHH is supportive of overall growth/viability of aggregates, but not essential for the induction of the adrenocortical lineage. These findings suggest that adrenocortical specification might be the default pathway of PIM lineage when inductive cues towards other lineages (e.g., nephrogenic lineage) are not provided. The elucidation of mechanisms by which NODAL/ACTIVIN or NOTCH dampen adrenocortical specification warrants further investigation.

Although prolonged floating culture of aggregates was hindered by decreased viability, subsequent ALI culture successfully induced further maturation of cells, which bore remarkable histologic and, ultrastructural similarities with FZ adrenocortical cells. Importantly, these cells produced substantial quantities of Δ5 steroids (i.e., DHEA, DHEA-S) and low/undetectable quantities of Δ4 steroids (e.g., cortisol, aldosterone), features characteristic of human fetal adrenocortical cells^1^. This is in stark contrast to earlier studies in which adrenocortical-like cells induced from hiPSCs/ESCs or multipotential stem cells primarily produced Δ4 steroids, which functionally closer to adult adrenocortical cells^29–31^. In contrast to our study in which NR5A1 (NT) was upregulated spontaneously, these prior studies over-expressed NR5A1, and it is therefore possible that steroidogenesis was induced directly by NR5A1 without the appropriate cellular or epigenetic context. Moreover, these prior studies also utilized agonists of cAMP/PKA signaling for induction of adrenocortical cells. Notably, in our study, adrenocortical lineages were induced in an ACTH/cAMP/PKA independent manner, which is more in accord with human adrenocortical development in vivo where ACTH is critical for growth but not for specification or initial development^1,32,33^.

Our transcriptome analyses revealed progressive and directional developmental trajectories of the adrenocortical lineage with remarkable similarity to those observed in vivo. Surprisingly, single cell transcriptome analyses also revealed the presence of minor cell types that could not be detected by bulk analyses, which included WT1^+^ nephrogenic cells and *RSPO3*^+^ capsule-like cells, both of which arose from the same progenitor as the adrenocortical lineage according to trajectory analyses (Fig. 5a). In mice, pre- and postnatal adrenal capsule surrounding the cortex produces the critical niche, which provides RSPO3 to subjacent adrenal cortex, thereby maintaining adrenocortical stem cells, that are in turn critical for homeostasis of the adrenal cortex and zonation^34^. Similar cell types were also identified in human fetal adrenals, but their origin was unknown^1,12^. Our transcriptomic comparison supports the notion that CapLCs, which closely resemble Cap in vivo, might originate from the PIM/CE similar to the adrenocortical lineage.

We also noted that during the trajectory towards FZLCs, there was a transient upregulation of DZ markers (e.g., *NOV*, *HOPX*)^12^. Accordingly, IF of ali28 revealed the presence of cells with DZ markers. Our recent scRNA-seq analysis of human fetal adrenals has recently demonstrated that FZ fate can be established either through direct differentiation from the AP (direct route) or indirectly through the DZ (indirect route)^12^. In the trajectory analyses in this study, DZLCs appear to locate in the midst of the trajectory leading to FZLCs, suggesting that FZLCs might be generated indirectly through DZLCs (Extended Data Fig. 6f). However, DZLCs were not identified as a distinct cell cluster, presumably due to the relatively small number of DZLCs captured in our scRNA-seq. Thus, the lineage relationship between FZLCs and DZLCs remains inconclusive, and it is still possible that some of the FZLCs might be induced directly. Interestingly, DZLCs preferentially locate at the periphery of the cluster of FZLCs where it is juxtaposed to mesenchymal cell types including the CapLCs. Thus, it is tempting to speculate that CapLC-provided instructive cues such as RSPO3, might potentiate canonical Wnt/β-catenin signaling to promote the DZ-like fate. The requirement for CapLCs in the formation of DZLCs warrants future investigation.

The first human identified with a *NR5A1* mutation had a heterozygous amino acid change (p.G35E) in the P-box (DNA-binding domain), resulting in primary adrenal insufficiency along with gonadal failure (classic phenotype)^35^. While a number of heterozygotes with mutations outside G35E have been described, and a handful of heterozygote mutants and a single hypomorphic homozygous mutant have been reported to show adrenal failure^36,37^, the rarity of mutations, varying clinical severity and effects of potential confounding factors (e.g., unrecognized [epi]mutations in regulatory regions, dominant negative effect) have made it difficult to determine the role of NR5A1 in human fetal adrenal development, posing significant clinical challenges in management and genetic counselling of patients and their family bearing NR5A1 mutations^38^. In this study, we revealed that homozygous null *NR5A1* mutants showed marked loss of cell viability and decreased steroidogenesis in the remaining cells, accompanied by downregulation of genes encoding steroidogenic proteins (e.g., *STAR*, *CYP11A1*, *CYP17A1*) (Fig. 7). Interestingly, mutant NT^+^ cells at fl22 exhibited substantial transcriptional similarities with wild-type NT^+^ cells at fl14, suggesting that NR5A1 is the major driver of the fl14-to-fl22 transition, in which marked upregulation of genes related to steroidogenesis normally occurs. Notably, some of the genes upregulated through the fl9-to-fl14 transition coinciding with the upregulation of NR5A1 (NT) were previously annotated as AP markers (e.g., *STRA6*, *RUNX1T1*, *GATA6*, *DLK1*). The initial increase in these markers and that of NR5A1 was not affected in the *NR5A1* mutant line, suggesting that there are yet unidentified mechanisms involved in activation of these early adrenocortical genes. Of note, upon ALI culture, NR5A1 mutant lines gradually reduce NT expression. This finding suggests that NR5A1 autoregulates its expression after its initial upregulation by as yet-to-be identified transcription factors. In support, NR5A1 binding motifs were identified in distal and proximal regulatory regions of the *NR5A1* loci. Future studies will systematically identify NR5A1-responsive genes and cis-regulatory mechanisms governing their expression through a multimodal sequencing approach to enhance our mechanistic understanding of how NR5A1 orchestrates FZ survival and steroidogenesis in human fetal adrenals. Such findings will not only allow us to provide a precise diagnosis for patients with NR5A1 mutations, but will identify novel gene regulatory networks, defects in which might lead to PAI or adrenocortical carcinoma^39,40^.

Based on the platform we developed, we envision several future directions. First, upstream genes that triggers expression of *NR5A1* and other early adrenocortical genes at fl14 could be explored individually by gene targeting or systematically by a functional genomics approach, offering a reverse genetics approach to predict the functional outcome of potentially hazardous mutations in human adrenal development. Second, this platform may also disclose novel information regarding adrenal androgen biosynthesis, which is unique to humans and non-human primates^41,42^, and insight into diseases associated with adrenal androgen excess (e.g., polycystic ovary syndrome)^43^. Third, further characterization and selective expansion of DZLCs might provide an opportunity to derive zona glomerulosa (zG) and fasciculata (zF), which originate from DZ perinatally^1,44^. However, this will require a further understanding of the developmental processes underlying differentiation of DZ and their postnatal derivatives, which might be informed by single cell analyses of human perinatal adrenal glands.

In summary, this in vitro human adrenocortical induction platform will serve as a critical foundation for understanding human adrenal development and its dysfunction, and will also serve as a stepping stone for cell replacement therapy for patients with adrenocortical dysfunction.

## METHODS

### Collection of human embryo samples

The fetal adrenal glands (9, 17, 19 wpf) and the heart at 9 wpf utilized for bulk RNA-seq or histologic analyses were obtained from donors undergoing elective abortion at the University of Pennsylvania, Fukuzumi Obstetrics and Gynecology Clinic. Embryo ages were determined by ultrasonographic measurement of the crown lump length or head circumference. All experimental procedures were approved by the Institutional Review boards at the University of Pennsylvania (#832470) and the Hokkaido University (19-066). The sex was determined with sex-specific PCR performed on genomic DNA with primers specific to the ZFX/ZFY loci^15^. The adrenal glands or the heart were submerged in RPMI-1640 medium and isolated from the surrounding connective or adipose tissues under a stereomicroscope.

### Feeder free culture of human iPSCs

Parental hiPSC lines, Penn123i-SV20 (male) and Penn067i-312-1 (female), were obtained from the iPSC core facility at the University of Pennsylvania^45^.1390G3 (female) was obtained from Dr. Masato Nakagawa at the Center for iPS Cell Research and Application, Kyoto University^46^. For maintenance of hiPSCs, cells were cultured on 6-well plates (Thermo Fisher Scientific) coated with iMatrix-511 Silk (Nacalai USA) in StemFit Basic04 medium (Ajinomoto) supplemented with 50 ng/mL basic FGF (Peprotech) or StemFit Basic04CT (complete type) medium at 37 °C under 5% CO2. For passaging or use for induction into adrenocortical lineages, hiPSC at day 6-7 after passaging were treated with a 1:1 mixture of TrypLE Select (Life Technologies) and 0.5 mM EDTA/phosphate-buffered saline (PBS) for 12-15 min at 37 °C to dissociate them into single cells. 10 μM ROCK inhibitor (Y-27632; Tocris) was supplemented in a culture medium 24 h after passaging hiPSCs. For single cell cloning of human iPSCs bearing WGNT or NT fluorescence reporters, human iPSCs were cultured in StemFit Basic03 medium supplemented with 50 ng/mL basic FGF.

### Generation of hiPSC lines bearing knock-in fluorescent reporter alleles

For a donor vector construction to generate the *WT1-p2A-EGFP* (WG) alleles, homology arms spanning the 3’ end of *WT1* loci (left arm:1157 bp; right arm: 1287 bp) were PCR-amplified from the genomic DNA of male hiPSCs (Penn123i-SV20) and were sub-cloned into the pCR2.1 vector using the TOPO TA cloning kit (Life Technologies). The *p2A-EGFP* fragment with the *PGK-Puro* cassette flanked by *loxP* sites was PCR-amplified from and inserted in-frame at the 3’-end of the *WT1* coding sequence using the GeneArt Seamless Cloning & Assembly Kit (Life Technologies). The *WT1* stop codon was removed to allow in-frame p2A-EGFP protein expression. Likewise, for donor vector construction to generate the *NR5A1-p2A-tdTomato* (NT) alleles, homology arms spanning the 3’ end of *NR5A1* loci were PCR-amplified from genomic DNA of male hiPSCs (585B1 8-6-8, a gift from Drs. Kotaro Sasaki and Mitinori Saitou, Kyoto University) and subcloned into the pCR2.1 vector^47^. The *p2A-tdTomato* fragment with the *PGK-neo* cassette flanked by the *loxP* site was also PCR-amplified and inserted in-frame at the 3’-end of the NR5A1 coding region. For the donor vector construction to establish *NR5A1^−/−^; NR5A1-p2A-tdTomato-polyA* alleles [NT knock-in, knock-out (NT-KIKO)], the genomic region spanning exons 2 and 3 of *NR5A1* loci from Penn123i-SV20 was first cloned in PCR2.1, which constituted a part of homology arms. This vector was linearized by inverse PCR skipping two zinc finger domains (critical domains for DNA-binding activities) constituting most of exons 2 and 3. This left only the first 42 bases of exon 2 and the last 38 bases of exon 3 in the homology arm (left arm: 892 bp; right arm: 1234 bp). Then, the *p2A-tdTomato-SV40 polyA* fragment with the *PGK-neo* cassette flanked by the *loxP* site was PCR-amplified and inserted in-frame to replace the zinc finger domains using a GeneArt Seamless Cloning & Assembly Kit. Finally, to reduce the random integration events, an *MC1-DT-A-polyA* cassette was PCR-amplified and inserted into the WG, NT or NT-KIKO donor vector, as described previously^48^.

Pairs of single-guide RNAs (sgRNAs) targeting the sequence at the 3’ end of *WT1* or *NR5A1*, exon 2 or 3 of *NR5A1* were designed using the Molecular Biology CRISPR design tool (Benchling). Pairs of sgRNA sequences were as follows: *WT1* 3’ end (5’-TCTGATGCATGTTGTGATGG-3’ and 5’-ACTCCAGCTGGCGCTTTGAG-3’); *NR5A1* 3’ end (5’-CTTGCAGCATTTCGATGAGC-3’ and 5’-CAGACTTGAGCCTGGGCCG-3’); *NR5A1* exon 2 (5’-AAGGTGTCCGGCTACCACTA-3’ and 5’- ACACCTTGTCCCCGCACACG −3’); *NR5A1* exon 3 (5’-CCGCTTCCAGAAATGCCTGA-3’ and 5’-TCTTGTCGATCTTGCAGCTC-3’). The sgRNAs were cloned into the *pX335-U6-Chimeric BB-CBh-hSpCas9n* (D10A) expression vector to generate the sgRNAs/Cas9n vector (a gift from Dr. Feng Zhang, Addgene, #42335; http://n2t.net/addgene:42335; RRID: Addgene_42335)^49^.

For generating hiPSCs bearing WGNT double fluorescent reporter alleles (WGNT SV20-211), the donor vectors (5 μg) and sgRNAs/Cas9n vectors (1 μg each) targeting *WT1* and *NR5A1* loci were simultaneously introduced into one million hiPSCs (Penn123i-SV20, male) by electroporation using a NEPA21 Type II Electroporator (Nepagene). After drug selection with puromycin and neomycin, cells were transfected with a plasmid expressing Cre recombinase (7.5 μg) to remove the *PGK-Puro* and *PGK-Neo* cassettes. These cells were plated into 96 well plates (1 cell/well) using the single-cell plating mode of FACSAria Fusion. After 10-12 days of culture using StemFit 03 medium in 96 well plate and 4-6 days in 12 well plates, cells were harvested; half of them were used for genotyping, and the other half were cryopreserved. To generate hiPSCs bearing an NT single fluorescence reporter (NT 312-2121 and NT 1390G3-2125), the donor and sgRNAs/Cas9n vectors targeting *NR5A1* loci were introduced into hiPSCs (Penn067i-312-1 or 1390G3, female) followed by neomycin selection. To generate KIKO hiPSCs (*WGNT-NR5A1^−/−^* SV20-211-19), hiPSCs (Penn123i-SV20) were transfected with the donor vector and sgRNAs/Cas9n targeting the 3’ end of *WT1* loci and subjected to puromycin selection. These cells were subsequently transfected with a donor vector and two pairs of sgRNAs/Cas9n targeting *NR5A1* exons 2 and 3, respectively, and subjected to neomycin selection. The cells were then transfected with a plasmid expressing Cre recombinase and subjected to single-cell plating using FACSAria Fusion. Clones bearing targeted alleles, random integration, or Cre recombination were determined by genotyping PCR using the primer pairs listed in Supplementary Table 6. Sequences of targeted alleles or non-targeted alleles were confirmed to be devoid of indels or other unexpected mutations by Sanger sequencing. Based on this screening, correctly targeted clones without random integration were selected. Thus, WGNT SV20-211 hiPSCs and WGNT SV20-KIKO19 were homozygous for WG and NT, NT 1390G3-2125 hiPSCs were homozygous for NT, and NT 312-2121 were heterozygous for NT. For all genetically modified cells, a master and secondary cell bank were generated in the following fashion. First, cell lines were expanded until there were sufficient cells to cryopreserve at least five vials as a master cell bank. Second, one vial from a master cell bank was thawed and expanded to create at least 20 cryovials as the secondary cell bank. Directed induction experiments were conducted using the secondary cell bank. We verified that all cell lines banked are mycoplasma free using MycoAlert mycoplasma detection assay (Cambrex). All experiments used passage number matched iPSCs (between p25 and p35). Cell lines used for experiments were not passaged more than six times after thawing to minimize the risk of culture-induced genetic changes.

### Karyotyping and G-band analyses

hiPSCs were incubated with 100 ng/ml of KaryoMAX Colcemid solution (Gibco) for 10 h in culture medium. Cells were dissociated using TrypLE Select and incubated with hypotonic solution (75 mM KCl) for 30 min at 37 °C. The cells fixed by Carnoy’s solution were dropped onto glass slides in a moisty chamber to prepare chromosomal spread. The number of chromosomes was counted using DAPI staining and the cell lines confirmed to be harboring 46 chromosomes were further analyzed for G-banding using Cell Line Genetics (Madison, WI).

### Induction of FZLCs though floating and ALI culture

To generate three-dimensional aggregates, hiPSCs were plated at 10,000 cells per well in V-bottom 96-well low-cell-binding plates (Greiner Bio-One) in DB27 medium (DMEM/F12 HEPES, 2% B27 supplement, 1% Glutamax, 1% insulin-transferrin-selenium [ITS-G], 1% MEM non-essential amino acids Solution, 90 μM 2-mercaptoethanol, 50 U/ml penicillin-streptomycin [all obtained from Thermo Fisher Scientific]), in the presence of 10 μM Y27632 (R&D Systems) and 2 ng/ml human BMP4 (R&D Systems). The plates were centrifugated at 210 g for 4 min to promote cell aggregation before incubation at 37 °C under 5% CO2 normoxic condition. After 24 h (at fl1), aggregates were transferred to U-bottom 96-well low-cell-binding plates (Greiner Bio-One) in a mesoderm-inducing medium containing 0.3 ng/ml BMP4 (R&D Systems) and 10 μM of CHIR99021 (R&D Systems). On fl3, half of the medium was replaced with fl1 medium containing 0.3 ng/ml BMP4, 10 μM of CHIR99021 and 10 μM Y-27632. On day 5, aggregates were transferred to MK10 medium (Minimum Essential Medium alpha [α-MEM, Thermo Fisher Scientific], 10% KSR (Thermo Fisher Scientific), 55 μM 2-mercaptoethanol, 100 U/ml penicillin/streptomycin (Thermo Fisher Scientific) containing 0.3 ng/ml BMP4, 2 μM SB-431542 (Sigma), 10 μM of CHIR99021 and 10 μM Y-27632. On fl7, aggregates were transfer to an MK10 medium containing 1 ng/ml BMP4, 30 μM SB-431542, 2 μM of CHIR99021, 0.1 μM retinoic acid, 50 ng/mL human sonic hedgehog/SHH (R&D Systems) and 10 μM Y-27632. On fl10, aggregates were transfer to MK10 medium containing 3 ng/ml BMP4, 30 μM SB-431542 (Sigma), 50 ng/mL human SHH, 10 μM of IWR-1 (Sigma), 2 μM SU-5402 (Tocris), and 10 μM DAPT (Tocris). On fl13, half of the medium were replaced with the same medium on fl10. On fl15, aggregates were transfer to MK10 medium containing, 2 μM SB-431542 (Sigma), 50 ng/mL SHH, 1 μM of IWR-1, 1 μM SU-5402, and 10 μM DAPT. On fl22, aggregates were transferred to BioCoat Collagen I Inserts with 0.4 μm PET membranes (Corning, 354444), which were subsequently placed on 24-well plates (Falcon) containing MK10 medium with 1 μM IWR-1, 5 μM DAPT, 50 ng/mL SHH, and 2 μM SU-5402. The inserts were placed in 24-well containing 200 μL of MK10 medium to equilibrate for 1 h prior to use. The inserts in 24-well plates were incubated at 37 °C and 5% to initiate ALI culture. The medium was changed every three days until harvesting aggregates for experiments.

Metanephric mesenchyme (MM) induction from hiPSCs (described in Figures S2A, C) was performed based on a previous study with minor modifications ^16^. Briefly, DB27 medium was used in all processes and the cytokine cocktail was as follows; [fl0; 0.5 ng/ml BMP4 and 10 μM Y-27632], [fl1; 1 ng/mL BMP4 and 10 μM of CHIR99021], [half medium change on fl03 and fl05; 1 ng/mL BMP4; 10 μM CHIR99021; 10 μM Y-27632], [fl7; 10 ng/ml activin A (R&D), 3 ng/ml BMP4, 3 μM CHIR99021, 0.1 μM retinoic acid, and 10 μM Y-27632], [fl9; 1 μM ChIR99021, 5 ng/ml human FGF9, and 10 μM Y-27632].

### Fluorescence-activated cell sorting (FACS)

Aggregates were dissociated using 0.1% trypsin/EDTA for 10-15 min at 37 °C with periodic pipetting. After the reaction was quenched by adding an equal volume of fetal bovine serum, cells were resuspended in FACS buffer (0.1% bovine serum albumin [BSA] in PBS). Cell suspensions were strained through 70-μm nylon cell strainers (Thermo Fisher Scientific) to remove clumps before sorting or analysis. To analyze/sort cells using a cell surface marker, the dissociated cells were stained with APC-conjugated anti-human DLK1 (R&D systems). Cells stained with the cell surface marker and those expressing NT were sorted using FACSAria Fusion (BD Biosciences) and were collected in 1.5-mL Eppendorf tubes or 15-mL Falcon tubes containing CELLOTION (Amsbio). All FACS data were collected using FACSDiva Software v 8.0.2 (BD Biosciences) and analyzed using FlowJo software (BD Biosciences).

### Quantitative RT-PCR

Cells were pelleted by centrifugation after isolation (500 g for 10 min) and subsequently lysed for RNA isolation using RNeasy Micro Kit (Qiagen). For some assays, cDNA synthesis was performed using Superscript III (Invitrogen) with oligo-dT primers and used for qPCR assays without amplification. In the remaining assays, cDNAs were synthesized and amplified using 1 ng of purified total RNA as described^47^. ERCC spike-in RNA sequences were added to the samples to estimate the transcript copy number per cell. Target cDNAs were quantified using PowerUp SYBR Green Master Mix (Applied Biosystems) with StepOnePlus (Thermo Fisher Scientific). The gene expression values were calculated using ΔCt (log2 scale) normalized to the average Ct values of *PPIA* and *ARBP*.

### Immunofluorescence analyses

The procedure was performed as described previously^12^. Briefly, aggregates were fixed with 4% paraformaldehyde (Sigma) in PBS for 2 h at room temperature, washed three times with PBS containing 0.2% Tween-20 (PBST) for 15 min, and successively immersed in 10% and 30% sucrose (Fisher Scientific) in PBS overnight at 4 °C. The samples were embedded in OCT (Fisher Scientific), snap-frozen with liquid nitrogen, and sectioned to 10-μm using a cryostat (Leica, CM1800). Sections were placed on Superfrost Microscope glass slides (Thermo Fisher Scientific) and stored at −80 °C until further analysis. The slides were air-dried, washed three times with PBS and incubated with a blocking solution (5% normal donkey serum in PBS containing 0.2% Tween-20) for 1 hr prior to primary antibody incubation. Several antibodies required antigen retrieval before the incubation. Slides were post-fixed with 10% buffered formalin (Fisher Healthcare) for 10 min at room temperature and washed with PBS three times for 10 min. Antigens were retrieved using HistoVT One (Nacalai USA) for 20 min at 70 °C and washed in PBS for 10 min at room temperature. Slides were subsequently incubated with primary antibodies in blocking solution for 2 h at room temperature, followed by washing with PBS six times (total of 2 h). Subsequently, slides were treated with secondary antibodies and 1 μg/ml DAPI (Thermo Fisher Scientific) in a blocking solution for 50 min at room temperature. Slides were washed with PBS six times and mounted in Vectashield mounting medium (Vector Laboratories) for confocal laser scanning microscopy analysis (Leica, SP5-FLIM inverted). Confocal images were processed using Leica LasX (version 3.7.2).

### IF analyses on paraffin sections

For IF analyses of fetal adrenal glands and aggregates during ALI culture, samples were fixed in 10% buffered formalin (Fisher Healthcare) with gentle rocking overnight at room temperature. After dehydration, tissues were embedded in paraffin, serially sectioned at 4-μm using a microtome (Thermo Scientific Microm™ HM325), and placed on Superfrost Microscope glass slides. Paraffin sections were then de-paraffinized using xylene. The procedure of IF was as previously described (Cheng et al., 2022). Briefly, slides were treated with HistVT One for 35 min at 90 °C, then 15 min at room temperature for antigen retrieval. The slides were incubated with primary antibodies in blocking buffer overnight at 4 °C. The slides were washed six times with PBS, followed by incubation with secondary antibodies in blocking buffer and 1 μg/ml DAPI (Thermo Fisher Scientific) in blocking solution for 50 min at room temperature. This step was followed by washing with PBS for six times before mounting in Vectashield mounting medium for confocal laser scanning microscopy.

### Antibodies

The primary antibodies were as follows: mouse anti-NR5A1 (R&D Systems, PP-N1665-0C; RRID: AB_2251509) , mouse anti-GATA4 (Santa Cruz Biotechnology, sc-25310; RRID:AB_2108747), goat anti-GATA4 (Santa Cruz Biotechnology, sc-1237, RRID:AB_2108747), rabbit anti-WT1 (Abcam, ab89901; RRID:AB_2043201), rabbit anti-CYP11A1 (ATLAS ANIBODIES, HPA016436; AB_1847423), rabbit anti-CYP17A1 (Abcam, ab134910; RRID: AB_2895598), rabbit anti-WT1 (Abcam, ab89901; RRID:AB_ 2043201), rabbit anti-SULT2A1 (Novus Biologicals, NBP2-49402; RRID: N.D.), rabbit anti-cleaved caspase-3 (Cell Signaling Technology, 9661; RRID: AB_2341188), mouse anti-human Pref-1/DLK1/FA1 APC-conjugated antibody (R&D Systems, FAB1144A; RRID:AB_2890004), and mouse anti-human CD10 (Biocare Medical, PM129AA, RRID; AB_10582488). The secondary antibodies were as follows: Alexa Fluor 488 conjugated donkey anti-rabbit IgG (Life Technologies, A21206; RRID:AB_2535792), Alexa Fluor 488 conjugated donkey anti-mouse IgG (Life Technologies, A21202; RRID: AB_141607), Alexa Fluor 568 conjugated donkey anti-mouse IgG (Life Technologies, A10037; RRID:AB_2534013), Alexa Fluor 568 conjugated donkey anti-rabbit IgG (Life Technologies, A10042; RRID:AB_2534017), Alexa Fluor 647 conjugated donkey anti-goat IgG (Life Technologies, A21447; RRID:AB_2535864), Alexa Fluor 647 conjugated donkey anti-rabbit IgG (Life Technologies, A31573; RRID:AB_2536183) and Alexa Fluor 647 conjugated donkey anti-mouse IgG (Life Technologies, A31571; RRID: AB_162542).

### In-situ hybridization (ISH) on paraffin sections

ISH analysis on formalin-fixed paraffin-embedded sections was performed as described ^12^. Briefly, the samples were hybridized with a ViewRNA ISH Tissue Assay Kit (Thermo Fisher Scientific) with gene-specific probe sets for human *NR5A1* (#VA1-3002602-VT), *APOA1* (#VA1-10349-VT), *STAR* (#VA1-3004623-VT), *NR0B1* (#VA1-3001628-VT), *NOV* (#VA1-3000396-VT), *RSPO3* (#VA1-13645-VT) and *PDGFRA* (#VA1-12826-VT). Probe sets against Bacillus S. *dapB* (#VF1-11712-VT) were used as the negative control. Sections deparaffinized by xylene were treated with pretreatment solution for 12 min at 90–95 °C, followed by a protease solution for 6 min and 30 sec at 40 °C. After fixation in 10% formaldehyde neutral buffer solution for 5 min, sections were hybridized with the ViewRNA type 1 probe set for 2 h at 40 °C. This step was followed by incubation with the preamplifier probe (25 min at 40 °C), the amplifier probe (15 min at 40 °C) and the Label Probe 1-AP (15 min at 40 °C). The sections were subsequently incubated with the AP enhancer for 5 min at room temperature, followed by development using FastRed Substrate Solution for 1 h at room temperature. The slides were counterstained with DAPI (1 μg/mL) in PBS for 1 h, then mounted in Vectashield mounting medium for confocal laser scanning microscopy.

### Measurement of steroid hormones

Medium for aggregates was refreshed three days before collection for steroid hormone measurement. In some experiments, substrates and various agonists or inhibitors were added when the medium was changed 3 days before collecting supernatants. After 72 h of incubation, supernatants were collected and stored at −20 °C until use. Commercially-available clinical-grade enzyme-linked immunosorbent assay (ELISA) kits (DRG International) were used to detect pregnenolone (EIA-4170), DHEA (EIA3415), and DHEA-S (EIA1562). All procedures were according to the manufacturer’s instructions.

Spike and recovery and precision experiments were conducted using supernatants obtained from 211 hiPSC-derived aggregates at ali21 as validation of the ELISA kit; %recovery was calculated using the following equation: (spiked sample concentration − sample concentration) *100 / spiked standard diluent concentration. Two concentrations of spiked standard diluent were used to calculate %recovery: pregnenolone (3.2 ng/ml and 12.8 ng/ml), DHEA (5 ng/ml and 15 ng/ml), and DHEA-S (2.5 μg/ml and 5 μg/ml). Four technical replicates were measured in a precision experiment. The optical density (OD) value of 450 nm (OD450 nm) was determined using Synergy 2 (BioTek) and processed using Gen5 software (BioTek). Steroid production levels were normalized to 1×10^3^ NT^+^ cell numbers in some experiments. Treatment effects of trophic stimulants were compared with the non-treated group (control), and inhibitors were compared with the LDL (50 μg/ml) + forskolin 1 μM group. Statistical significance was analyzed using one-way ANOVA for repeated measures followed by Dunnett’s multiple comparison test.

### Steroid quantification by liquid chromatography-tandem mass spectometry (LC-MS/MS)

Progesterone, 17-OH, aldosterone, 11-deoxycorticosterone (DOC), androstenedione (A4), 11β-hydroxyandrostenedione (11OHA4), and 11-ketoandrostenedione (11KA4), were measured using LC-MS/MS. Briefly, the culture medium (100 μl) was combined with internal standards, diluted with deionized water, and loaded onto a supported liquid extraction cartridge (Isolute, Biotage, Charlotte, NC). After 5 minutes, steroids were eluted twice with 0.7 mL methyl-tert-butyl ether, dried under vacuum (Savant, Thermo Fisher), and reconstituted in 0.1 mL 40:60 methanol:water. LC-MS/MS was performed as previously described ^50^. The steroids were quantified in dynamic multiple reaction monitoring modes. The lower limit of quantification for each steroid was estimated from the signal-to-noise ratio of pooled samples and extrapolated to a concentration that achieved a signal-to-noise ratio of 3.

Creative Proteomics measured progesterone, cortisol, and testosterone (T). Briefly, the culture medium (150 μl) was extracted with solid phase extraction and dried under the nitrogen. Samples were then reconstituted with 100 μl methanol (50 ng/mL progesterone-d9 as internal standard for LC-MS/MS analysis). AB SCIEX Qtrap 5500 tandem mass spectrometer connected to a Waters Acquity UPLC was used for the analysis. A Waters Acquity UPLC BEH C18 column (2.1×100 mm, 1.7 μm) was used, and the column temperature was set to 40 °C, and the injection volume was 6 μl. The mobile phase consisted of 0.1% formic acid 2 mM ammonium fluoride solution (phase A), and acetonitrile (phase B). The mass spectrometer was operated in positive mode using an ESI source.

### Bulk RNA-seq library preparation

The fetal adrenal glands and the heart, hiPSCs, and floating and ALI cultured aggregates were used for bulk RNA-sequencing. According to the manufacturer’s instructions, the fetal tissues were dissociated using a Multi Tissue Dissociation Kit 1 (Milteny Biotech). After tissues were minced into <1-mm fragments using scissors, samples were collected in 15-ml Falcon tubes and mixed with dissociation solution (enzyme mix mixed with serum-free DMEM). Then, samples were incubated at 37 °C for 30 minutes in a water bath with pipetting every 10 minutes. At the end of incubation, fetal bovine serum was added (15% of total volume), and cell suspensions were strained through 70-μm cell strainers. Cells were washed once in PBS by centrifugation at 300 g for 5 minutes, and subsequently treated with Red Blood Cell Lysis Solution (Miltenyi Biotec) for 2 minutes at room temperature. After adding 3 ml DMEM, cells were centrifuged at 300 g for 5 min and resuspended in FACS buffer before cell number was counted using trypan blue exclusion. In some samples (fetal adrenal glands at 17 and 19 wpf), cells were subsequently stained with APC-anti-CD10 antibody, and CD10 negative cells were sorted using FACSAria Fusion. Cells were pelleted by centrifugation at 300 g for 5 min and pellets were snap-frozen in liquid nitrogen and stored at −80 °C until RNA isolation.

Dissociation of hiPSC-derived aggregates was as described previously^48^. The following samples were prepared: fl0 hiPSC (WGNT SV20-211 [male, n =2]), aggregates at fl9 (whole WG^+^ fraction, WGNT SV20-211 [n=2]), fl14 (NT^+^ fraction, WGNT SV20-211 [n=2]), fl22 (NT^+^ fraction, WGNT SV20-211 [n=2]), fl22 (NT^+^ fraction, NT 312-2121 [female, n=2]), fl22 (NT^+^ fraction, WGNT SV20-KIKO19 [male, n=2]), ali21 (NT^+^ fraction, WGNT SV20-211 [n=1]), ali21 (NT^+^ fraction, NT 1390G-2125 [n=1]) and ali 28 (NT^+^ fraction, WGNT SV20-211 [n=1]). RNA was extracted using an RNeasy Micro Kit (Qiagen) and RNA quality was evaluated using High Sensitivity RNA Screen Tape on an Agilent 4200 TapeStation. The cDNA library was prepared using Clontech SMART-Seq® HT Plus Kit (PN R400749) according to the manufacturer’s protocol. AMPure XP beads were used for library cleanup. 75-base pair reads were sequenced on an Illumina NextSeq 500 machine according to the manufacturer’s protocol.

### Mapping reads and data analysis for bulk RNA-seq

Raw sequencing data were demultiplexed as Fastq files using bcl2fastq2 (v2.20.0.422). Barcode and adaptors were removed using Trimmomatic (v0.32). Fastq files were mapped to a UCSC human reference genome (GRH38) using STAR (2.7.1a). A raw gene count table was generated using featurecounts. DEGs were analyzed using edgeR (v3.36.0) with log2 fold change above 2, P-value < 0.05, and FDR < 0.05; log2(FPKM+1) was used to make scatterplots.

### 10x Genomics scRNA-seq library preparation

WGNT SV20-211 hiPSC-derived aggregates were collected for scRNA-seq analyses: aggregates at fl19 (FACS-sorted whole WG^+^ fraction [n= 1] or NT^+^ fraction [n=1]), fl22 (NT^+^ fraction [n=1]), ali21 (whole WG+ fraction [n=1]) and ali28 (NT^+^ fraction [n =1]). Aggregates were dissociated in 0.1% Trypsin/EDTA for 10-15 min at 37 °C with periodic pipetting. After the reaction was quenched by adding an equal volume of fetal bovine serum, cells were resuspended in FACS buffer (0.1% BSA in PBS). Cell suspensions were strained through a 70-μm nylon cell strainer (Thermo Fisher Scientific) to remove cell clumps before use. All samples were stained with trypan blue and confirmed to be >85% viable. Cells were loaded into a Chromium microfluidic chip B using the Chromium Single Cell 3ʼ Reagent Kit (v3 chemistry) and Chromium Controller (10X Genomics) to generate gel bead emulsions (GEMs). Gelbead emulsion-RT was performed using a C1000 Touch Thermal Cycler equipped with a deep-well head (Bio-Rad). Subsequent cDNA amplification and library construction steps were performed according to the manufacturer’s instruction. Libraries were sequenced using a NextSeq 500/500 high output kit v2 (150 cycles) (FC-404-2002) on an Illumina NextSeq 550 sequencer.

### Mapping reads and data analyses for scRNA-seq

Raw data were demultiplexed with the mkfastq command in cellranger (V5.0.1) to generate Fastq files. Trimmed sequence files were mapped to the reference genome for humans (GRCh38) provided by 10x Genomics. Read counts were obtained from outputs from Cell Ranger.

Secondary data analyses were performed in R (v.4.1.0) with Seurat (v.4.0). UMI count tables were first loaded into R using the Read10X function, and Seurat objects were built from each sample. Cells with fewer than 200 genes, an aberrantly high gene count above 7000, or a percentage of total mitochondrial genes >20% were filtered out. Of the 30,319 cells for which transcriptomes were available, 23,570 cells passed the quality-control dataset filters and were used in downstream analysis. We detected 3572 median genes/cell at a mean sequencing depth of 44,499 reads/cell (Extended Data Fig. 6a). Samples were combined, and the effects of mitochondrial genes and cell cycle genes were regressed out with SCTransform during normalization in Seurat; then, gene counts were scaled by 10000 and natural log normalized. Mitochondrial genes and cell cycle genes were excluded during cell clustering, dimensional reduction and trajectory analysis. Cells were clustered according to Seurat’s shared nearest neighbor modularity optimization-based clustering algorithm. Clusters were annotated based on previously characterized marker gene expression, and cluster annotation was generated for downstream analyses. Dimensional reduction was performed with the top 3000 highly variable genes and the first 30 principal components with Seurat. Differentially expressed genes (DEGs) across different clusters calculated using Seurat findallmarkers with thresholds of an average log2-fold change above 0.25 and p < 0.01. Developmental trajectories of cells were simulated using PHATE. Trajectory principal lines were fitted with ElPiGraph.R. Pseudotime was calculated using ElPiGraph.R. The cell cycle was analyzed using CellCycleScoring in Seurat. DEGs between two groups in scatterplots were identified using edgeR 3.34.1 by applying a quasi-likelihood approach and the fraction of detected genes per cell as covariates. The DEGs were defined as those with FDR <0.01, p < 0.01 and a log2-fold-change >0.25. For pseudo-bulk analyses, DEGs were defined as those with FDR <0.01, p <0.01, and a log2-fold-change >1. Data were visualized using ggplot2 and pheatmap. Genes in the heatmap were hierarchically clustered according to Euclidean distance, scaled by row and then visualized with pheatmap. Gene ontology enrichment was analyzed using DAVID v6.8.

### Library preparation, mapping reads and data analyses for scATAC-seq

FACS-sorted cells of floating day 9 (fl9) and day 22 (fl22) samples [WT1 positive population from fl9 (211), n=1; NR5A1 positive population from fl22 (211), n=1] were used for scATAC-seq library preparation with a Chromium Single Cell ATAC Reagent Kits (v1.1 Chemistry). All cells preparation, nuclear extraction, subsequent cDNA amplification and library construction steps were performed according to the manufacturer’s instruction. Libraries were sequenced using NovaSeq 6000 v1.5 reagents kit on an Illumina NovaSeq 6000 sequencer.

The scATAC-seq mapping to the human reference genome was provided by 10X genomics using the cellranger-atac-2.0.0 pipeline. The downstream analysis was performed using Signac (v1.6.0). Generally, initial QC metrics were calculated using Signac, and peak_region_fragments > 3000, peak_region_fragments < 100000, pct_reads_in_peaks > 40, blacklist_ratio < 0.025, nucleosome_signal < 4, and TSS.enrichment > 2 were used for filtration of low-quality cells. Clustering and the remaining analyses followed default parameters. Briefly, term frequency-inverse document frequency (TF-IDF) normalization was performed. The top 95% of most frequently observed features (peaks) were used for dimension reduction and clustering. Iterative latent semantic indexing (LSI) was used for dimensional reduction. Cell clusters were identified using FindClusters in Signac with smart local moving algorithm. The gene activity score was calculated based on the number of fragments for each cell that map to gene and promoter regions (gene body plus 2 KB upstream of TSS). The motif activity score was calculated using chromVAR (v1.16.0) and the JASPAR CORE database (v2020). Transcription factor footprinting was performed using Footprint in Signac. Trajectory was constructed using PHATE with the same peak matrix for clustering. Trajectory principal lines were fitted with ElPiGraph.R. Pseudotime was calculated with ElPiGraph.R. Pseudo-bulk analysis was performed using HOMER (v4.6). Peaks were called by MACS2 and annotated by ChIPpeakAnno (v3.28.1). Genome coverage was generated by DeepTools (v3.5.1) and visualized in the IGV genome browser (v 2.12.3).

### Data and software availability

Accession numbers generated in this study are GSE201794 (10x Chromium scRNA-seq data for hiPSC-derived aggregates), GSE201793 (10X Chromium scATAC-seq data for hiPSC-derived aggregates) and GSE201792 (bulk RNA-seq for hiPSC-derived aggregates and the fetal samples). The codes used for pseudotime analysis are available at GitHub repository).

## ACKNOWLEDGMENTS

We thank L. King for carefully reviewing the manuscript and providing insightful comments. We thank the Women’s Health and Clinical Research Center and the Tumor Tissue and Biospecimen Bank at the University of Pennsylvania for human sample collection and T. Moriwaki for making histologic sections and assisting immunohistochemistry. We thank members of Sasaki lab for the discussion of this study.

This work was supported in part by the Open Philanthropy funds from the Silicon Valley Community Foundation (2019-197906) and the Good Ventures Foundation (10080664) to K.S and P30-ES013508 (NIEHS) to Center of Excellence in Environmental Toxicology at the University of Pennsylvania.

## AUTHOR CONTRIBUTIONS

K.Sasaki conceived and supervised the project and designed the overall experiments. K.Sasaki, J.F.S, K.C., M.M. and Y.Sakata wrote the manuscript. M.M., Y.Sakata, K.C., and K.Sasaki conducted the overall experiments and analyzed the data. K.C., contributed to the analyses of RNA-seq and scATAC-seq data. M.M. and Y.Seita. contributed to the processing of human embryo materials. A.D., T.M.P, K.Shishikura, and R.J.A contributed to the LC-MS/MS analyses and data interpretation. W.Y generated parental hiPSCs.

## SUPPLEMENTARY TABLES: see separate Excel documents

**Supplementary Table 1.** DEGs identified from a pairwise comparison between samples at different time points during FZLC induction.

**Supplementary Table 2.** DEGs identified from a multi-group comparison between hiPSCs, aggregates at fl09-14, fl22, and ALI culture.

**Supplementary Table 3.** DEGs identified from a pairwise comparison between AP and PIM/ECE from cynomolgus monkey embryos.

**Supplementary Table 4.** DEGs identified from a multi-group comparison between cell types in scRNA-seq analysis.

**Supplementary Table 5.** DEGs identified from a pairwise comparison between wild-type and mutant aggregates.

**Supplementary Table 6.** Primers used in this study

