## Supplemental figures for "Reconstitution of human adrenocortical specification and steroidogenesis using induced pluripotent stem cells"

### Extended Data Fig. 1

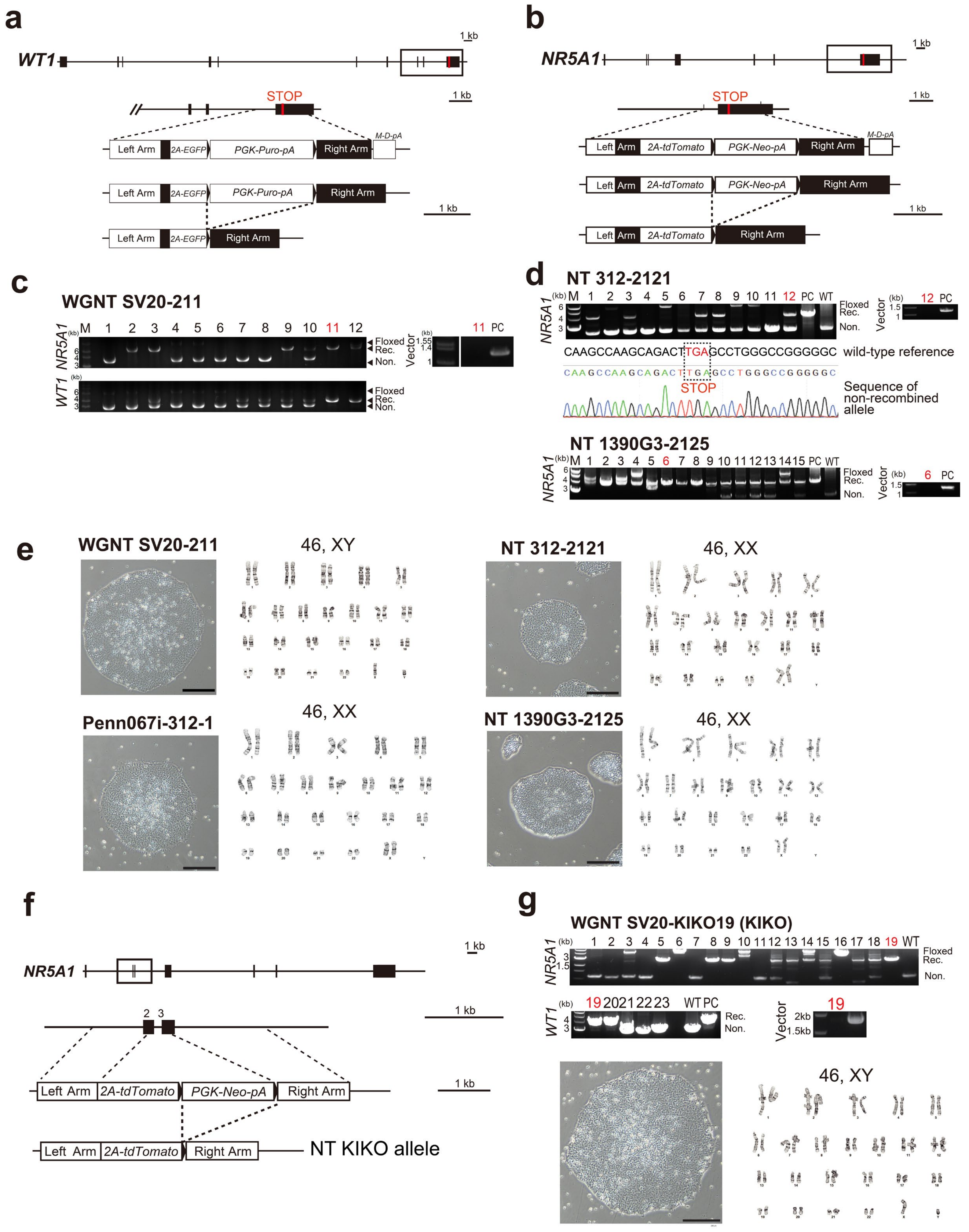

**Extended Data Fig. 1. The generation of hiPSCs bearing *WGNT* or *NT* fluorescence reporter alleles or *NR5A1* mutant alleles.** (**a**) Schematic illustration of the human *WT1* locus and the targeting construct for generating *WT1-p2A-EG-FP* alleles (WG). Black boxes indicate exons. (**b**) Schematic illustration of the human *NR5A1* locus and the targeting construct for generating *NR5A1-p2A-tdTomato* alleles (NT). Black boxes indicate exons. (**c**) Genotyping PCR to screen for clones bearing WG (top left) and NT (bottom left). Note that clone 11 (designated as WGNT SV20-211) bears WT and NT biallelically, and does not show random integration of the targeting vectors (top right), Floxed, floxed by loxP; Rec., recombination by Cre; Non., non-targeted. (**d**) Genotyping PCR to screen for clones bearing NT. Penn067i-312-1 and 1390G3 are used as parental hiPSCs. Note that clone 12 (top, designated as NT 312-2121) bears monoallelic NT, whereas the other allele is free of indels as confirmed by Sanger sequencing. Clone 6 (bottom, designated as NT 1390G3-2125) bears biallelic NT. Random integration of the targeting vectors is not observed in either clone. (**e**) Phase-contrast images and karyotype analysis of indicated hiPSCs. All four clones bear normal karyotypes (46, XY, or XX). Bar, 250 μm. (**f**) Schematic illustration of the human *NR5A1* locus and the targeting construct for generating *NR5A1-/-; NR5A1-p2A-tdTomato-polyA* alleles (NT-KIKO). Black boxes indicate exons. (**g**) Genotyping PCR to screen for clones bearing NT-KIKO and WG. Penn123i-SV20 is used as a parental hiPSCs. Note that clone 19 (designated as WGNT SV20-KIKO19) bears NT-KIKO and WG biallelically and is free of random integration of the targeting vector (top). A phase-contrast image (bottom left) and karyotype analysis of WGNT SV20-KIKO19 hiPSC, showing a normal karyotype (46, XY) (bottom right). Bar, 250 μ m.

### Extended Data Fig. 2

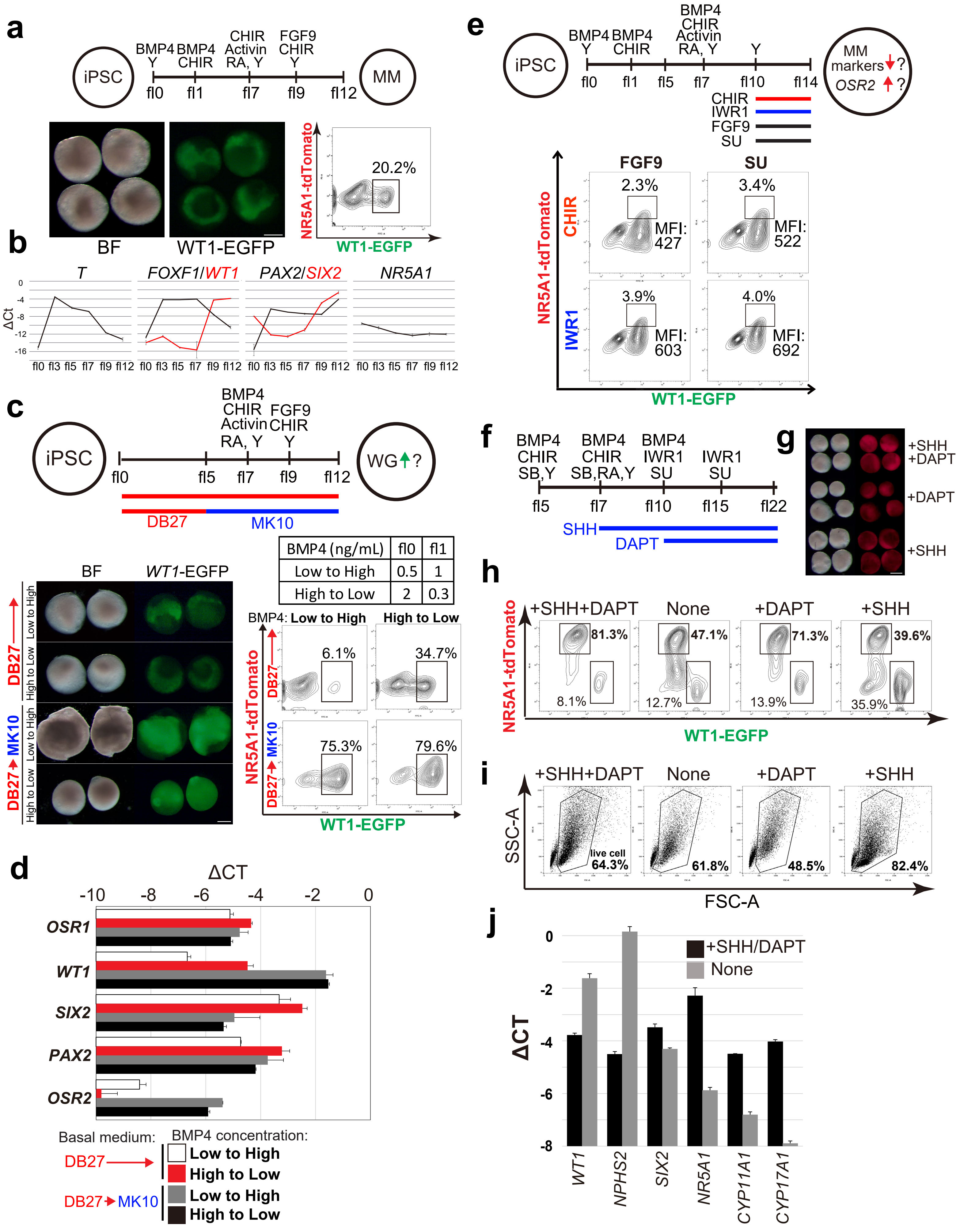

**Extended Data Fig. 2. Optimization of floating culture to establish NT$^+$ early adrenocortical lineage from hiPSCs.**
(**a**) Outline of the metanephric mesenchyme (MM) induction from hiPSCs based on a previous study with minor modifications (top, also see METHODS). Bright-field (BF) and fluorescence (WG) images (bottom left) and FACS analysis for WG expression (bottom right) of aggregates at fl12. CHIR, CHIR99021; RA, retinoic acid; Y, Y-27632. (**b**) The quantification of gene expression levels of indicated markers measured by qPCR during induction of MM as outlined in (**a**). (**c**) Experimental outline to increase the induction efficiency of WG$^+$ (top). Basal medium (DB27 [f0-12] versus combination of DB27 [fl0-5] and MK10 [fl5-12]) and BMP4 concentration during fl0-1 (shown in chart) are optimized. DB27, DMEM/F-12 medium supplemented with B27; MK10, MEM-α medium supplemented with 10% KSR. Outcomes were evaluated by BF and fluorescence (WG) images (left), and FACS analyses (right) at fl12. (**d**) qPCR quantification of key gene expression of bulk aggregates generated in (**c**). For each gene examined, the ΔCt from the average Ct values of the two independent housekeeping genes *ARBP* and *PPIA* (set as 0) were calculated and plotted. The average value from two independent experiments is shown on the log2 scale, with SDs. (**e**) FACS analyses to evaluate the effect of Wnt signaling (IWR1 [10 μM] or CHIR99021 [1 μM]) and FDF9 signaling (FGF9 [5 ng/ml] or SU-5402 [2 μM]) during fl10-14 (top). (**f**) Experimental scheme for evaluation of the effect of SHH (50 ng/ml) and DAPT (10 μM) during fl7-22. SB, SB-431542; SHH, human sonic hedgehog; SU, SU-5402. (**g**) Bright-field (BF) and NT images of aggregates at fl22 induced by indicated conditions as in (**f**). Bar, 500 μm. (**h**) FACS analyses of aggregates at fl22 as in (**f, g**). (**i**) Percentage of living cell as in (**h**). (**j**) The qPCR quantification of gene expression of bulk aggregates at fl22 treated with or without SHH/DAPT as in (**f**).

### Extended Data Fig. 3

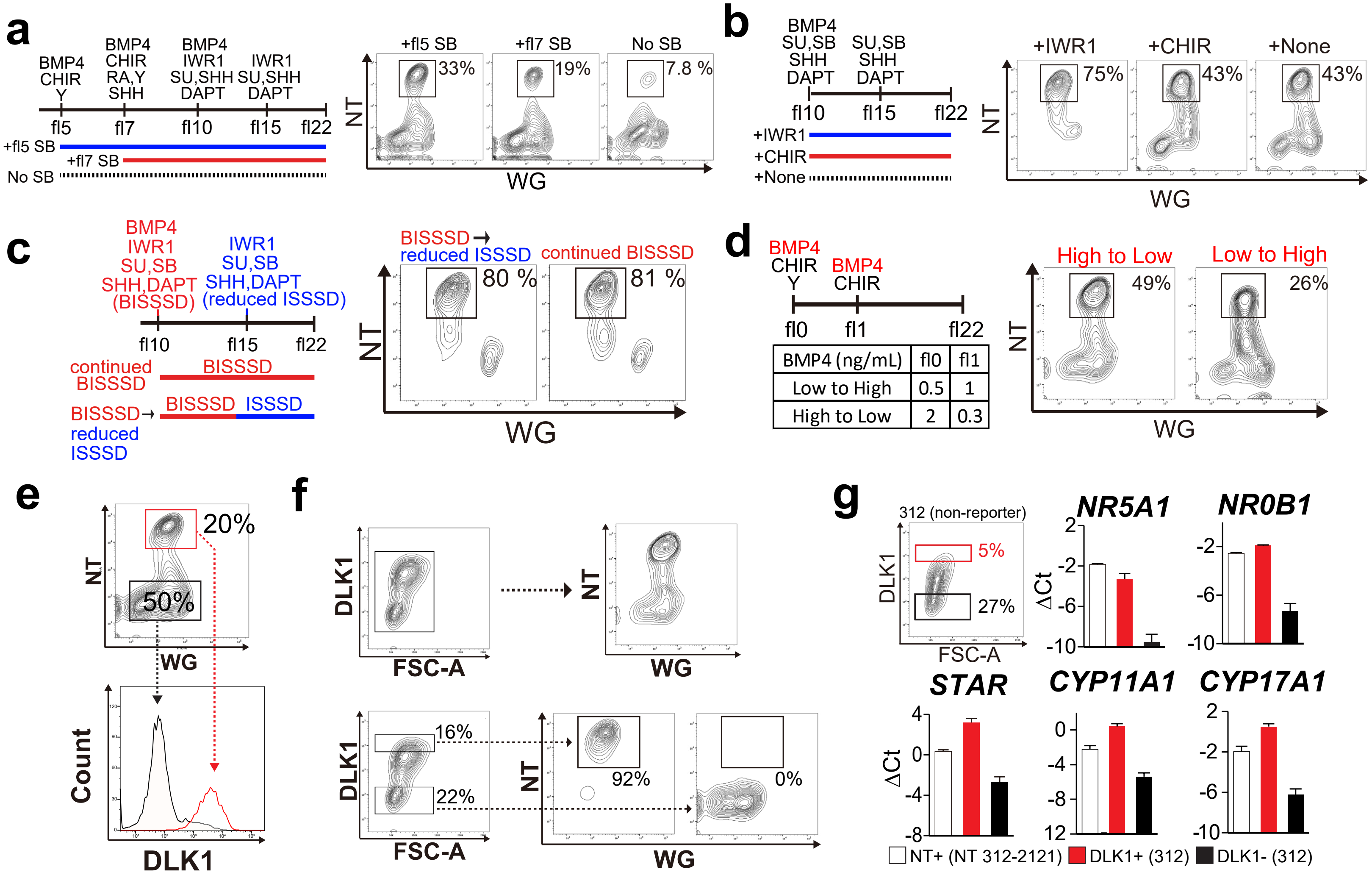


**Extended Data Fig. 3. Optimization of floating culture and identification of DLK1 as a surface marker of adrenocortical cells at fl22.** (**a**) FACS analysis to evaluate the effect of SB-431542 on induction of NT$^{+}$ cells added during fl5-22 or fl7-22 or not added. The remaining factors added are the same as in Fig. 1b. (**b**) Effect of IWR1 or CHIR on induction of NT+ cells during fl10-22. (**c**) Effect of different culture condition during fl10-22 on induction of NT$^{+}$ cells. Continued cultures with BISSSD (fl10-22) or cultures switched to ISSSD at fl15 are compared. BISSD contains BMP4, IWR1, SU, SB, SHH, and DAPT; ISSSD contains IWR1, SU, SB, SHH and DAPT. The remaining factors are the same as in Fig. 1b. (**d**) Effect of BMP4 during fl0-1 on induction of NT$^{+}$ cells. The remaining factors are the same as in Fig. 1b. (**e**) DLK1 expression in NT$^{+}$ or NT$^{-}$ cells from fl22 aggregates derived from 211 hiPSCs assessed by FACS. (**f**) WGNT expression in whole aggregates or cells gated on DLK1$^{+}$ or DLK1$^{-}$ fraction as assessed by FACS. 211 hiPSCs are used for induction. (**g**) FACS-sorting of DLK1$^{+}$ and DLK1$^{-}$ cells from aggregates at fl22 derived from non-reporter Penn067i-312-1 (312) hiPSCs. These cells, with sorted NT$^{+}$ cells from an NT 312-2121 reporter line, were evaluated for key gene expression by qPCR.

### Extended Data Fig. 4

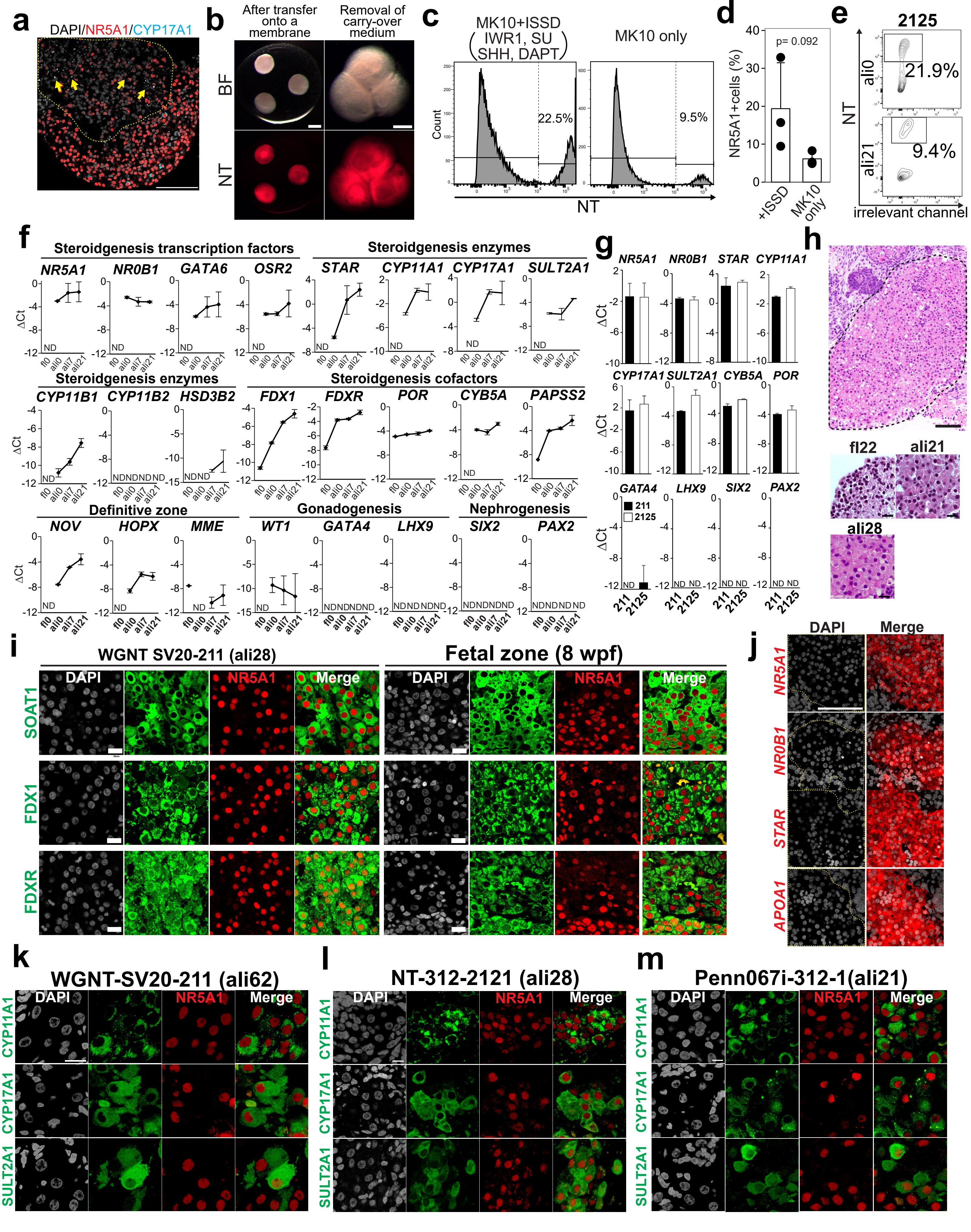

**Extended Data Fig. 4. Optimization of ALI culture and characterization of FZLC.** (**a**) Merged IF images of the fl24 for DAPI (white), NR5A1 (red) and CYP17A1 (cyan). Dotted lines and yellow arrows indicate dead cells. Bar, 100 μm. (**b**) Bright-field (top) and fluorescence images (bottom) of fl21 aggregates after transfer onto a membrane (left). The carry-over medium is subsequently removed, resulting in the merging of three aggregates into one aggregate (right). (**c**) FACS histogram of ali21 aggregates cultured with ISSD in MK10 medium (left) or MK10 only (right) for NT expression. (**d**) Proportion of $NT^+$ cells in ali21 aggregates as in (**c**). Means ± standard deviation (n = 3). (**e**) FACs plots of ali0 and 21 aggregates derived from NT 1390G3-2125 hiPSCs for NT expression. (**f**) qPCR quantification of key genes of FACS-sorted $NT^+$ cells derived from 211 hiPSCs at the indicated time points. N.D., not detected. (**g**) qPCR quantification of key genes of FACS-sorted $NT^+$ cells derived from 211 or 2125 hiPSCs at ali21. (**h**) H&E images of aggregates at ali28 at low magnification (top) and those at fl22, ali21 and ali28 at high magnification. Cells are derived from 211 hiPSCs. Bar, 100 μ m (top) and 20 μm (bottom). (**i**) IF images of ali28 aggregates derived from 211 hiPSCs or human embryo at 8 wpf, for indicated markers (green) co-stained with NR5A1 (red) and DAPI (white). Merged images are shown on the right. Bar, 20 μm. (**j**) In situ hybridization (ISH) of ali28 aggregates (derived from 211 hiPSCs) for the indicated genes (red) merged with DAPI (white). Dotted lines encircle the FZLCs. Bar, 100 μm. (**k**) IF images of ali62 aggregates derived from 211 hiPSCs for CYP11A1, CYP17A1 or SULT2A1 (green) co-stained with NR5A1 (red) and DAPI (white). Merges of NR5A1 and indicated marker are shown on the right. Bar, 10 μm. (**l**) IF images of ali28 aggregates derived from NT 312-2121 hiPSCs for indicated markers as in (**k**). Bar, 10 μm. (**m**) IF images of ali21 aggregates derived from Penn067i-312-1 hiPSCs for indicated markers as in (**k**). Bar, 10 μm.

### Extended Data Fig. 5

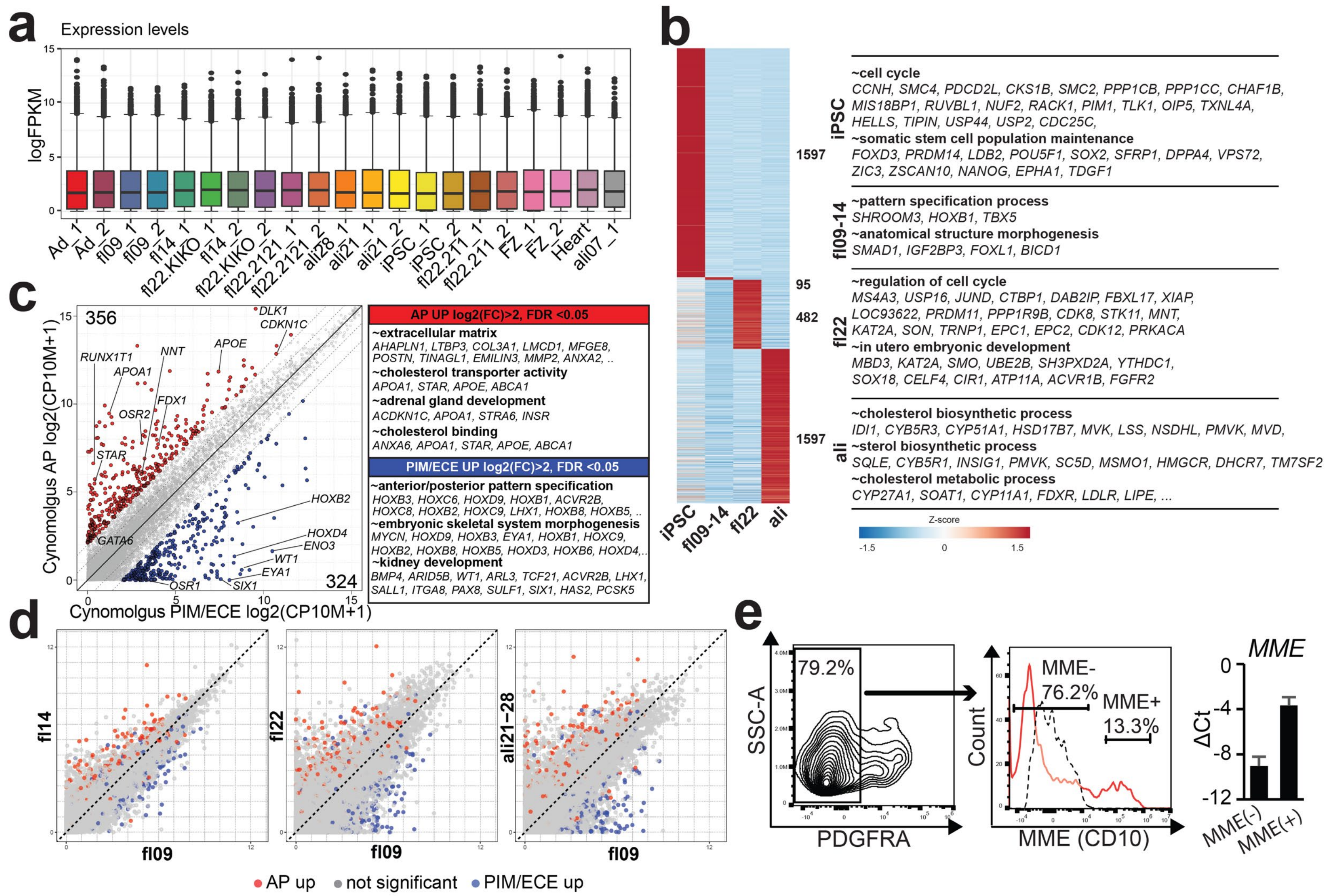


**Extended Data Fig. 5. Bulk transcriptome analyses to characterize FZLCs.** (**a**) Overall transcriptome expression levels among all bulk RNA-seq datasets. (**b**) Heatmap showing the averaged expression pattern of DEGs identified from a multi-group comparison between hiPSC, aggregates at fl09-14, fl22 and ALI culture, sorted based on the dendrogram clustering in Fig. 4a. Up-regulated genes with log-fold change above 0.5, $p < 0.05$ and FDR < 0.05 are identified. Enriched GO terms among DEGs are shown on the right. (**c**) (left) Scatter plot compares the averaged gene expression values between AP and PIM/ECE in cynomolgus monkey embryos 15. DEGs are identified with log-fold change > 2, FDR < 0.05. (right) GO analyses of the DEGs. Representative genes in each GO category are shown. (**d**) DEGs identified in (**c**) projected on scatter plots comparing averaged expression values between in vitro samples at the indicated stages (defined in Fig. 4a, c). (**e**) Isolation of $MME^+$ or $MME^-$ cells from $PDGFRA^-$ fraction of a 19 wpf human fetal adrenal gland using FACS (left) and qPCR quantification of *MME* expression of sorted cells. PDGFRA and MME are the adrenal capsule and definitive zone (DZ) markers, respectively. Accordingly, isolated $MME^-PDGFRA^-$ cells enriched with FZ were used for bulk RNA-seq in Fig. 4e.

### Extended Data Fig. 6

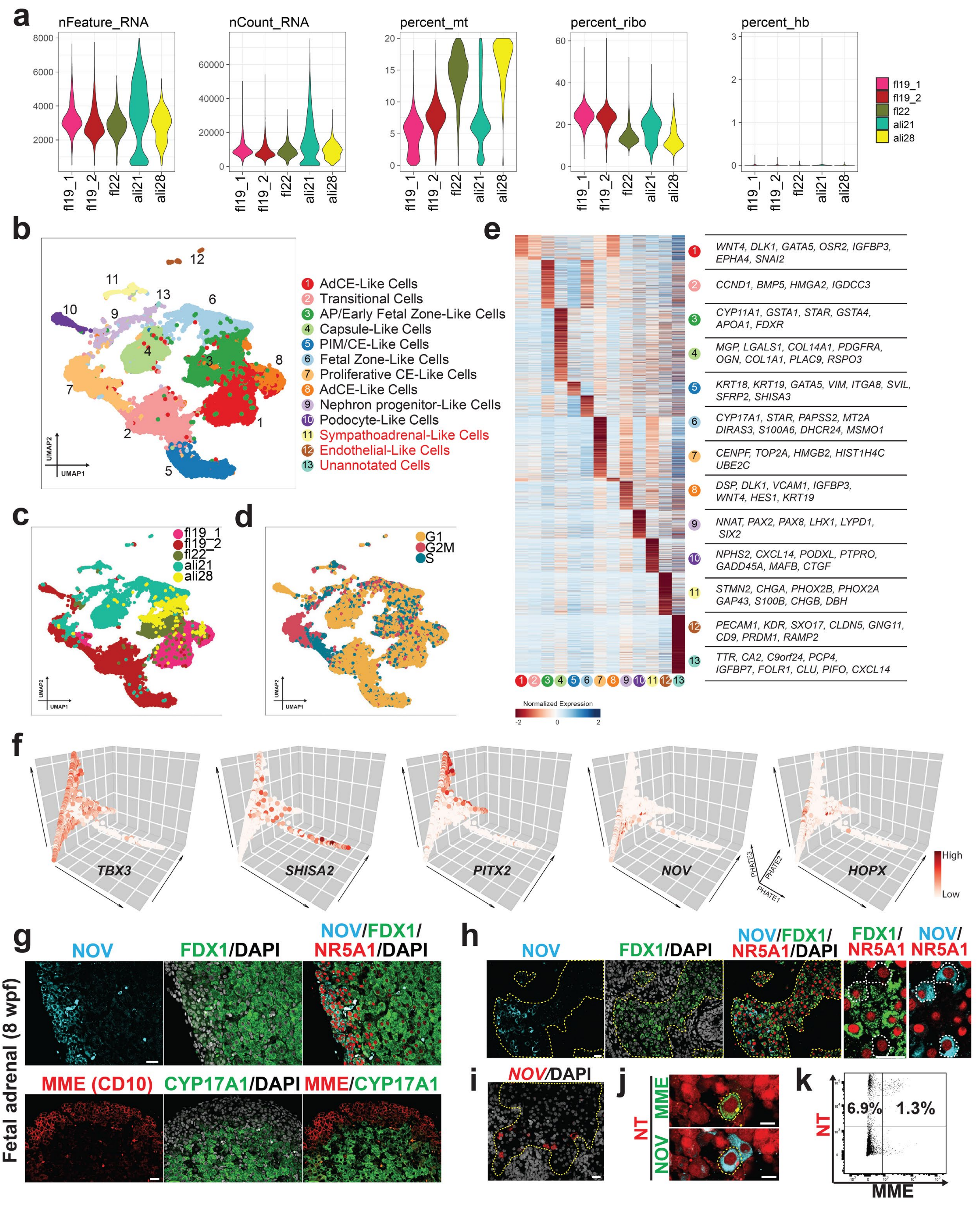

**Extended Data Fig. 6. scRNA-seq analyses to characterize FZLCs.** (**a**) Quality control metrics and associated plots of scRNA-seq. (**b-d**) UMAP plot of all cells based on computationally aggregated scRNA-seq data obtained from five different samples. Cells are colored according to clusters identified by Seurat clustering analysis (**b**), sample origin (**c**) or cell cycle status (**d**). Cells were annotated based on markers characteristic of indicated cell types, except cluster 13, which is not annotated. (**e**) Heatmap showing the averaged expression pattern of the DEGs identified from a multi-group comparison between cell types defined in (**b**) (log fold change > 0.25, $p < 0.01$). Log-normalized expression value is scaled by row with Z-score transformation. (**f**) Selected marker gene expression for PIM/CE (*TBX3*, *SHISA2*, *PITX2*) or definitive zone (*NOV*, *HOPX*) projected on PHATE embedding as shown in Fig. 5a. Color denotes the expression level. (**g**) IF of a human fetal adrenal gland (8wpf) for NOV (cyan), FDX1 (green), NR5A1 (red), and DAPI (white) with their merged images (top) or MME (red), CYP17A1 (green), and DAPI (white) with their merged images (bottom). NOV and MME represent DZ markers, and FDX1 and CYP17A1 represent FZ markers. Bar, 20 µm. (**h**) IF of an ali28 aggregate for indicated markers. Note that $NR5A1^{+}NOV^{+}FDX1^{weak+}$ DZ-like cells are preferentially localized at the periphery of $FDX1^{+}$ FZLCs. (**i**) ISH of an ali28 aggregate for *NOV* merged with DAPI. Bar, 20 µm. (**j**) IF of ali21 aggregates for MME (green) or NOV (cyan) merged with NT (red), highlighting some $NOV^{+}$ cells that also express MME. Bar, 10 µm. (**k**) FACS plot of ali21 aggregates for NT and MME (CD10) showing a small portion of $MME^{+}$ cells among $NT^{+}$ cells.

### Extended Data Fig. 7

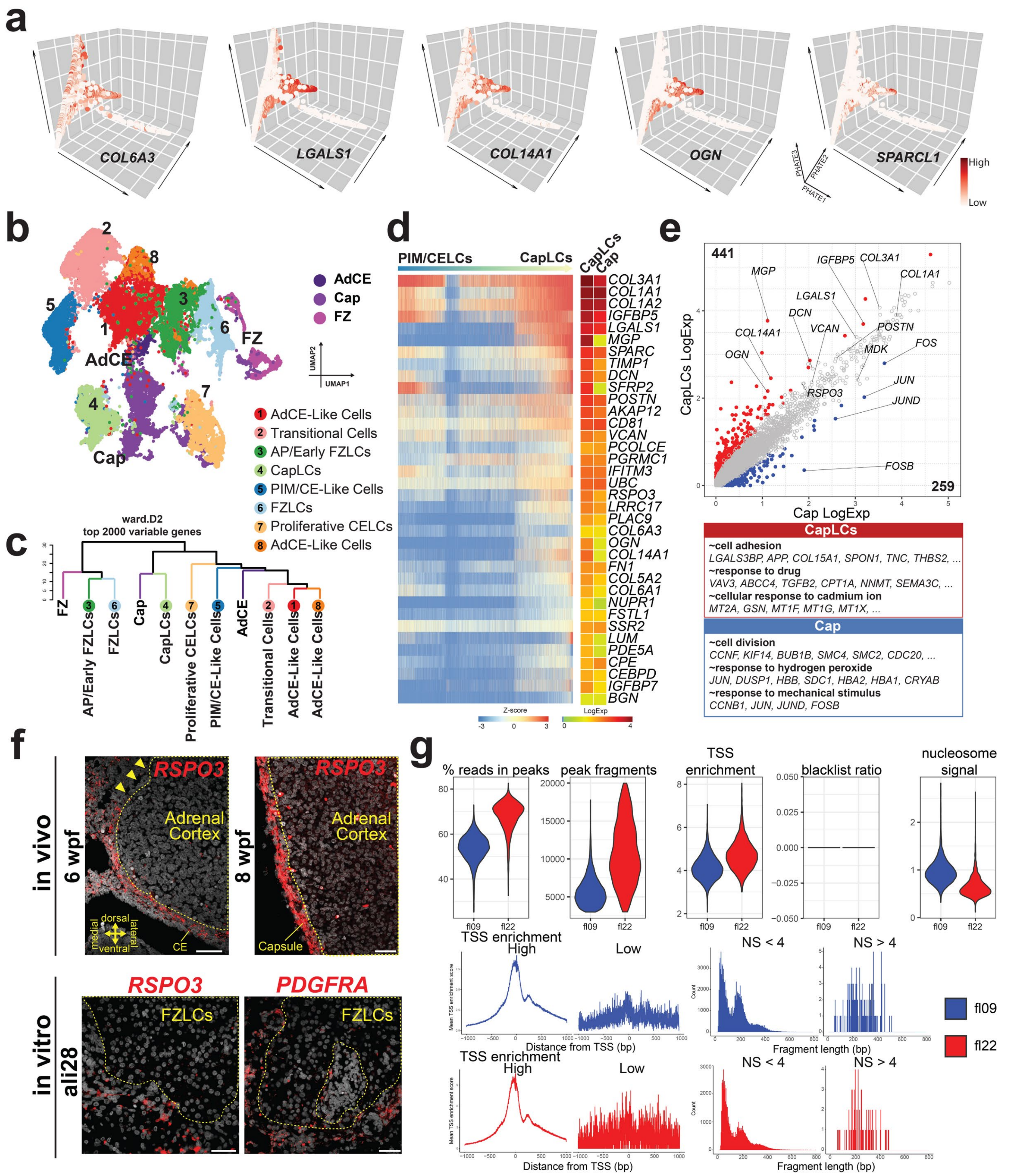

**Extended Data Fig. 7. Characterization of capsule-like cells by scRNA-seq and QC analysis of scATAC-seq.** (**a**) Expression of the adrenal capsule markers projected on PHATE trajectory defined in Fig. 5a. Markers are selected based on DEGs for the adrenal capsule cells (Cap) from our previous study12. (**b**) UMAP embedding showing the integration of in vivo (AdCE, Cap, FZ) and in vitro derived cells (clusters 1-8). Cell clusters 1–8 are defined in Fig. 5a and Extended Data Fig. 6b. In vivo cells are derived from 4–8 wpf human embryos and defined in our previous report[12]. (**c**) Dendrogram showing hierarchical cluster analysis of in vivo and in vitro derived cells. Cell clusters are ordered by ward.D2 using the top 2000 variable genes. (**d**) (left) Heatmap showing dynamic gene expression during the CapLC specification from PIM/CELCs. Single cells along the lineage trajectory of CapLCs are ordered by pseudotime as defined in Fig. 5c. The color indicates the Z-score of log-scaled expression. (right) Heatmap showing averaged gene expression values of CapLCs and Cap, and color indicates the average of log normalized expression. (**e**) Scatter plot comparison of pseudo-bulk transcriptomes between CapLCs and Cap. DEGs (log-fold change > 2, $p < 0.05$ and FDR < 0.05) are highlighted in colors; 441 genes were higher in CapLCs (red) and 259 genes were higher in Cap (blue). Key genes are annotated and representative genes and GO enrichments for DEGs are shown at the bottom. (**f**) ISH of the fetal adrenal glands at 6 wpf (top left) and 8 wpf (top right) for *RSPO3* or of ali28 aggregates derived from 211 hiPSCs for *RSPO3* (bottom left) or *PDGFRA* (bottom right). Adrenal cortex (adrenal primordium, AP) (top) or FZLCs are outlined by yellow dotted lines. Arrowheads denote the dorsomedial aspect of the AP that is devoid of Cap. (**g**) Quality-control metrics of scATAC-seq. TSS, transcription start site; NS, nucleosome.
